# The ligand-mediated affinity of escort proteins determines the directionality of lipophilic cargo transport

**DOI:** 10.1101/415984

**Authors:** Yi-Yun Cheng, Yun-Fang Huang, Hsin-Hui Lin, Wun-Shaing Wayne Chang, Ping-Chiang Lyu

## Abstract

Intracellular cargo transport is a highly dynamic process. In eukaryotic cells, the uptake and release of lipophilic ligands are executed by escort proteins. However, how these carriers control the directionality of cargo trafficking remains unclear. Here, we have elucidated the unliganded structure of an archetypal fatty acid-binding protein (FABP) and found that it possesses stronger binding affinity than its liganded counterpart towards empty nanodiscs. Titrating unliganded FABP and nanodiscs with long-chain fatty acids (LCFAs) rescued the broadening of FABP cross-peak intensities in HSQC spectra due to decreased protein-membrane interaction. Crystallographic studies revealed that the tails of bound LCFAs obstructed the charged interfaces of the FABP–nanodisc complexes. We conclude that the lipophilic ligands, by taking advantage of escort proteins with high conformational homogeneity and nanodiscs as the third interaction partner involved in this transport study, participate directly in the control of their own transportation in an irreversible, unidirectional fashion.

## Introduction

Intracellular cargo transport is the cornerstone of many biological processes, including cellular metabolism, energy homeostasis, neuron development and organelle function. In eukaryotic cells, a number of escort proteins have evolved to shuttle lipophilic cargos from external or internal environments to distinct subcellular compartments (1-3). Among these escort proteins, the cytosolic fatty acid-binding proteins (FABPs) are the archetypal lipid chaperones for long-chain fatty acids (LCFAs) and certain other energy-rich lipophilic ligands (4-7). The FABP carriage system is essential because 1) it minimizes the potential toxicity of free non-esterified LCFAs; 2) it facilitates the rapid cellular uptake and regulatable transport of LCFAs in concert with metabolic demands; and 3) it permits LCFAs to traverse intracellular space to various sites of utilization, e.g. to the endoplasmic reticulum for transacylation and to the nucleus for transcriptional regulation (8, 9). Upon binding to LCFAs, FABPs themselves may enter the nucleus to activate several transcription factors such as PPARs (peroxisome proliferator-activated receptors) and ERRs (oestrogen-related receptors) for downstream cellular signalling, metabolic reactions and immune responses (10-12).

Considerable evidence suggests that once absorbed by cell membrane phospholipids, LCFAs are sequestered and delivered by FABPs (13, 14) in a manner probably similar to the substrate-binding proteins (15). Despite substantial efforts to determine the key structural factors contributing to lipophilic cargo binding (16-18), the precise mode of how LCFAs are directionally transported by FABPs from a specific donor membrane to a target acceptor membrane or protein remain largely in dispute (2). One major reason for this ambiguity is inadequate research on the role of the phospholipid membrane in cargo trafficking, in that the uptake and release of lipophilic ligands depend not only on the ligand competition between escort proteins and ligand donors/acceptors but also on the selective affinities of unliganded and liganded escort proteins towards ligand donors/acceptors. Recent NMR and protein engineering studies revealed that the helical portal motif of FABPs for ligand entry was capable of reversibly transferring LCFAs across the plasma and intracellular membranes (19-22). However, owing to the uncertain conformational homogeneity (23, 24) and the lack of appropriate biomembrane mimetics in past research (25, 26), it is still unclear whether lipophilic ligands can be reversibly transported and how escort proteins control the directionality of cargo trafficking.

In this article, we utilized *Drosophila* FABP as a model system to assess the hypothesis of a directionality-controlled mechanism in escort proteins. An improved purification strategy was provided to separate the liganded-like and unliganded forms of FABP. The latter exhibited very high conformational homogeneity, allowing us to verify how different forms of FABP interact differently with lipid bilayer membranes. By employing membrane mimetic nanodiscs for NMR experiments, we were able to increase the tumbling rates of FABP–nanodisc complexes to reveal the residue-based membrane-interacting region of escort protein. Our structural analyses demonstrated that unliganded FABP has a much stronger binding affinity than its liganded counterpart to empty nanodiscs. Crystallographic studies showed that the LCFAs participate in their own transportation by disturbing the charged interfaces between FABPs and lipid membranes with aliphatic tails. Together, these results clearly suggest that the membrane-binding capacity of escort proteins is regulated by their target ligands, ultimately facilitating irreversible unidirectional lipophilic cargo transfer.

## Results

### Escort proteins with high conformational homogeneity are important for determining the regulatory mechanism of membrane-associated cargo transport

The escort protein represented herein is a *Drosophila* brain-type FABP (dFABP) that exists mainly in glial cells and localizes in both the cytoplasm and the nucleus (27, 28). Its movement from the cytoplasm to the nucleus in response to diurnal regulation and memory consolidation raises the question of how ligands trigger dFABP function. Although earlier biochemical assays of FABPs tolerated the minor heterogeneity in assessing the nature of their ligand binding (29, 30), we believed that studying escort proteins with high conformational homogeneity would illustrate the apparently dynamic lipid transport process. To clarify whether insufficient conformational homogeneity may impede further research on the interaction between FABPs and membrane mimetics, we compared the two apo forms of dFABP isolated by the existing method (denoted as the urea-treated method) and by the newly improved two-stage purification procedure with the use of the reagent GuHCl to induce partial unfolding (denoted as the GuHCl-purification method, see ***Supp. Fig. S1***). The results showed that the secondary structures of the two apo forms of dFABP with different conformational homogeneities varied significantly upon heating (***Fig. 1***). The impact of undesired conformational heterogeneity in dFABP on biophysical analysis was observed in the initial far-UV circular dichroism (CD) spectrum. In contrast to the irreversible profile observed for urea-treated dFABP, there exists a partially reversible profile for GuHCl-purified dFABP after the heating process (***Fig. 1A and B***). In addition, the melting curves obtained at 215 nm revealed that the two apo forms of dFABP behave differently upon interaction with the ligand oleic acid (OA; ***Supp. Fig. S2A***). This result implies that the secondary structures of the urea-treated sample were altered by the heat-induced precipitation of some nonprotein contaminants (***Supp. Fig. S3B and E***) that were absent in the GuHCl-purified sample (***Supp. Fig. S2D, S3F***). The undesired contaminants seem to have high molecular weights, as demonstrated by size-exclusion chromatography (denoted as P1 and A in ***Supp. Fig. S3B***). Despite the small change upon heating observed for the secondary structures of the apo and holo forms in GuHCl-purified dFABP, the melting temperatures measured through intrinsic fluorescence showed that both samples obtained from the two purification methods underwent tertiary structure denaturation (***Supp. Fig. S2C***).

**Figure 1.**
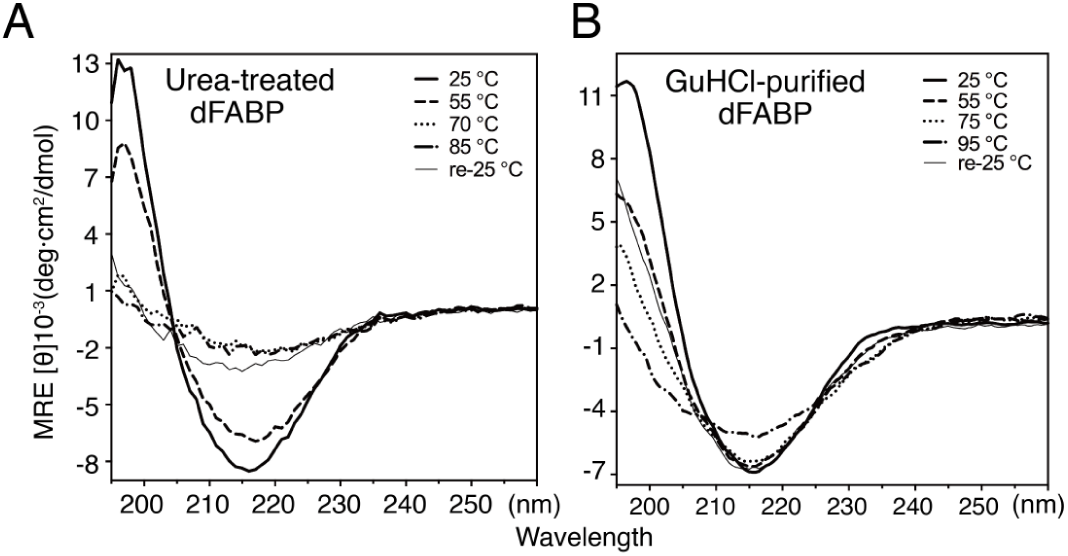
Spectroscopic analysis of urea-treated and GuHCl-purified dFABP. The urea-treated **(A)** and GuHCl-purified **(B)** samples were subjected to far-UV CD spectroscopy for secondary thermal stability analysis. Notably, the reversed 25 °C spectrum (denoted re-25 °C and shown as the thinnest line) in the temperature profile of the urea-treated dFABP was found to be irreversible at 215 nm due to protein precipitation ***(Supp. Fig. S2)***.

### Improved conformational homogeneity of dFABP benefits the study of membrane interaction by increasing the proportion of the apo-conformer

The pronounced difference observed in the melting curves of CD spectra convinced us to study the ligand binding based on the dFABP structure with different conformational homogeneities. The HSQC spectrum of the OA complex, holo-dFABP, was assigned (BMRB code 27112, ***Supp. Table S11***) and used as a template spectrum to manually assist the assignments of conformer-A and conformer-B’ (a conformer similar to the holo form) in the spectrum of apo-dFABP (BMRB code 27113, ***Supp. Table S12***). Conformer-B’ was identified as a liganded-like conformation. The cross-peaks in HSQC spectra indicated a higher proportion of conformer-A in the GuHCl-purified dFABP sample than in the urea-treated sample (***Fig. 2A and B***). Representative residues of conformer-A and conformer-B’ showed that the urea-treated apo form of dFABP might be a liganded-like dFABP based on the proportion of conformer-A, which was 3-fold less than that in the GuHCl-purified dFABP on average (***Fig. 2C and D***). Superimposition of the initial and final HSQC spectra upon OA titration revealed that the increasing intensity of conformer-B’ in urea-treated dFABP was difficult for comparison to the stoichiometry measured in biochemical assays (***Supp. Fig. S4D***). Although the disappearance of signals in conformer-A was similar to the results observed in GuHCl-purified dFABP (***Supp. Fig. S9***), the larger proportion of conformer-B’ in urea-treated dFABP proved to be problematic for subsequent structural analyses on membrane interaction.

**Figure 2.**
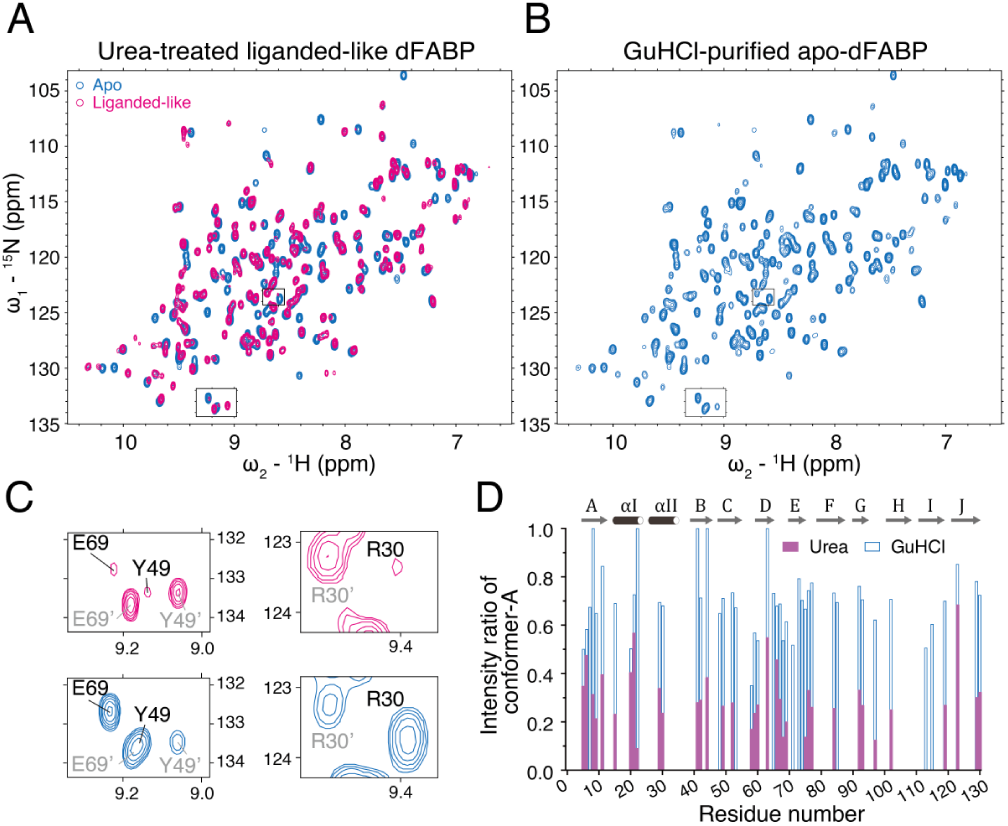
Conformational subpopulations of apo-dFABP determined by HSQC NMR. **(A)** The HSQC spectrum of urea-treated liganded-like dFABP is shown in magenta and is superimposed on that of GuHCl-purified dFABP (light blue). Representative residues for conformer comparison are marked using black boxes. **(B)** The HSQC spectrum of GuHCl-purified apo-dFABP is shown in light blue. Representative residues for conformer comparison are marked using black boxes. **(C)** Representative cross-peaks are enlarged to indicate the different conformational subpopulations in the apo forms of dFABP from the two purification strategies. The residues labelled in gray with the prime symbol represent conformer-B’, which is overwhelmingly present in urea-treated liganded-like dFABP, and the dispersion resembles that of OA-liganded holo-dFABP. **(D)** The intensity ratio of conformer-A reveals that the proportion of the apo form is approximately 3 times higher in GuHCl-purified dFABP than in urea-treated dFABP.

### Crystallographic structures of dFABP elucidate different potentials for membrane interaction with minimal protein conformational changes

The apo forms of both urea-treated and GuHCl-purified dFABP were crystallized, and their structures were obtained and identified as 5GGE and 5GKB, respectively (***Supp. Table S13***). Unexpectedly, the density maps for the ligand-binding cavity of urea-treated dFABP were incomplete, such that we were uncertain of the liganded state and therefore termed it the liganded-like structure (***Fig. 3A***). Since the region was relatively empty in GuHCl-purified dFABP, containing only three molecules of water, this structure was denoted as apo-dFABP (***Fig. 3B***). The crystal structures presented herein support the importance of conformational homogeneity through the subtle dissimilarities observed between two structures at similar resolution, with different temperature factors observed in the corresponding hypothesized portal (*α*II, loops of *β*C-D and *β*E-*β* and gap regions (*β*D-E). The temperature factor-derived atom distributions of *α*II and the *β*C-D loop in the portal region of apo-dFABP were more compact than those in liganded-like dFABP, and those of *β*E-F in the gap region were expanded during crystallization (***Fig. 3, Supp. Fig. S10***). The larger atom distributions of liganded-like dFABP in the portal region may have resulted from the use of ambiguous restraints for the aliphatic tail of the endogenous unknown ligand, while the internal hydration of water molecules within this region of apo-dFABP may result in more compact atom distributions (31). As shown in the 5GGE liganded-like dFABP structure, crystallization might preserve the slow-moving (gap region *β*E-*F*) and promiscuous binding (portal region *α*II) sites where diverse positions occur.

**Figure 3.**
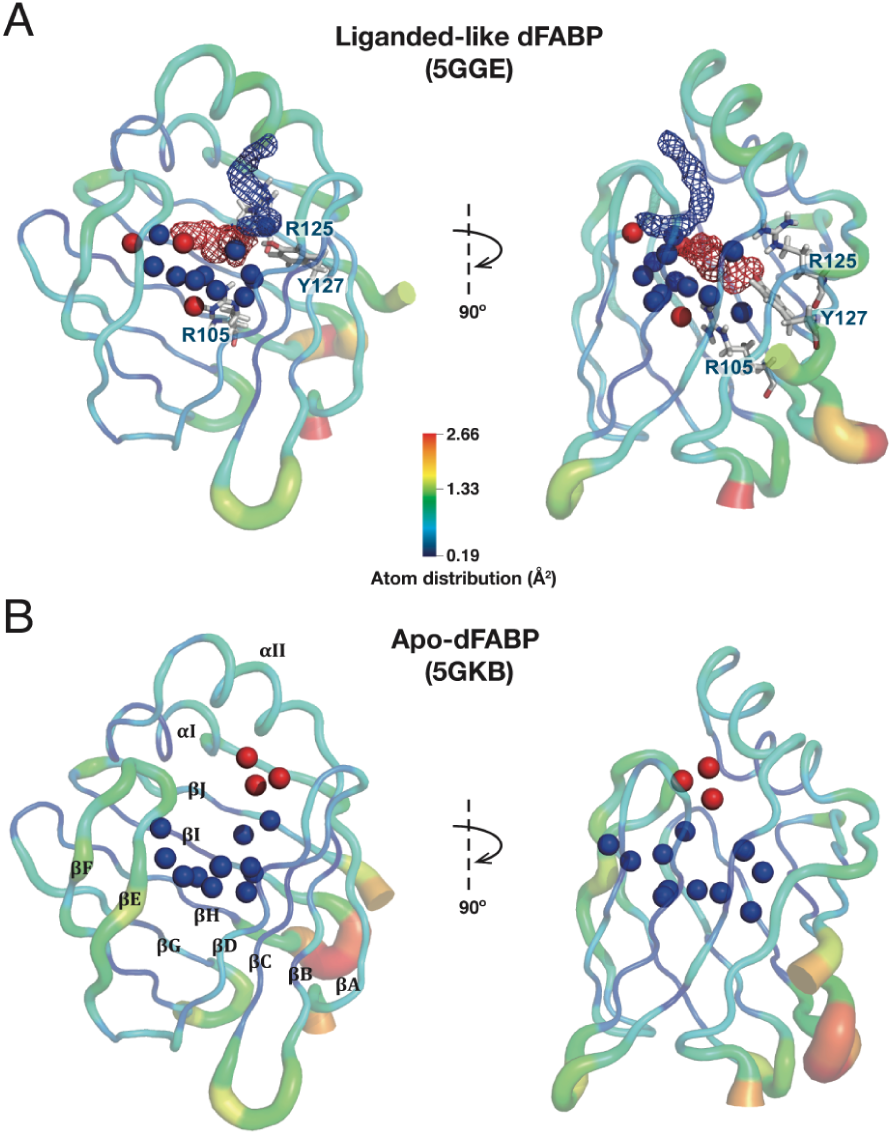
Distribution of water molecules in the apo structures of dFABP. **(A)** Fourteen water molecules within the ligand binding cavity are defined by PyMOL and shown as spheres. Eleven in similar locations are colored in blue. However, 3 water molecules (shown in red) were observed in different locations in the urea-treated liganded-like dFABP (PDB ID: 5GGE) and the GuHCl-purified apo-dFABP (PDB ID: 5GKB) structures. Both dFABP crystal structures are depicted with the true atom distribution derived from temperature factors, shown in a sausage representation. The discontinuously incomplete electron density maps are shown as red (lower) and blue (upper) meshes. The three residues R105, R125 and Y127, shown as CPK sticks, are within 3.5 Å of the lower part of the unknown ligand-like map and are conserved in FABPs due to their interaction with the carboxylate head of LCFAs. **(B)** The upper part of the cavity in the GuHCl-purified apo-dFABP is filled with 3 water molecules (shown in red) that are located in a different region from those in liganded-like dFABP. Two *α*-helices are nominated as *α*I and *α*II, and the ten *β*-strands are designated as *β*A-*β*J.

The altered distribution of ordered waters in liganded-like dFABP is worth comparing to that of apo-dFABP. Despite the presence of eleven water molecules within the binding cavity (blue) in both structures, three water molecules (red) were positioned differently (***Fig. 3***). Because of the existence of an unknown ligand, the three differently positioned water molecules were buried in the lower part of the cavity in the liganded-like dFABP but in the upper part of the cavity in apo-dFABP. In contrast to the possibility that the unknown ligand occupied the space in the upper cavity in the liganded-like dFABP, the allosteric ordered waters associated with apo-dFABP may be related to the rearrangement of the hydrogen networks occurring among the surrounding residues, as explained based on a prior molecular dynamics simulation of human heart-type FABP (32). Residues that are potentially involved in networks with a possible carboxylate group of the unknown ligand (within 3.5 Å) at the bottom of the ligand-binding cavity are also conserved in previous holo-FABP complex structures (***Fig. 3A, 5C***). Accordingly, the distribution of these water molecules ratherthan the unknown ligand near the upper cavity region of apo-dFABP is consistent with the claim of higher conformational homogeneity in the GuHCl-purified dFABP sample.

### Membrane interaction reveals a ligand triggering its own unidirectional transport through modulating the dissociation of apo-dFABP from membranes

The GuHCl purification of dFABP enabled a study on the role of conformational relationship in protein–membrane interactions. We first obtained conventional HSQC spectra to observe the discrepancies between the apo and holo forms of GuHCl-purified dFABP through titration with nanodiscs (***Fig. 4A and C***). Next, transverse relaxation-optimized spectroscopy (TROSY) spectra of apo- and holo-dFABP bound to the 1,2-dioleoyl-sn-glycero-3-phospho-(1’-rac-glycerol) (sodium salt) /1,2-dioleoyl-sn-glycero-3-phosphocholine (DOPG/DOPC) lipid nanodiscs (lipid NDs) at a 1:1 molar ratio were acquired to enhance the signal disappearance by virtue of the TROSY effect (***Fig. 4B and D, Supp. Fig. S5A, Supp. Table S14***). Contrary to our expectations, after the addition of the nanodiscs, holo-dFABP exhibited few changes from the previously acquired spectrum (***Fig. 4A***). In contrast, the signals of apo-dFABP in both the HSQC and TROSY spectra were clearly broadened after the addition of nanodiscs, by an order of magnitude after TROSY spectrum, due to a decrease in the tumbling rate (***Fig. 4B and C***). This result suggested that apo-rather than holo-dFABP interacts with membrane mimetic nanodiscs. The scale of peak broadening in apo-dFABP in the presence of lipid NDs was plotted as the fraction of attenuation and then superimposed on the tertiary structure of apo-dFABP in a sausage model (***Fig. 4D***).

**Figure 4.**
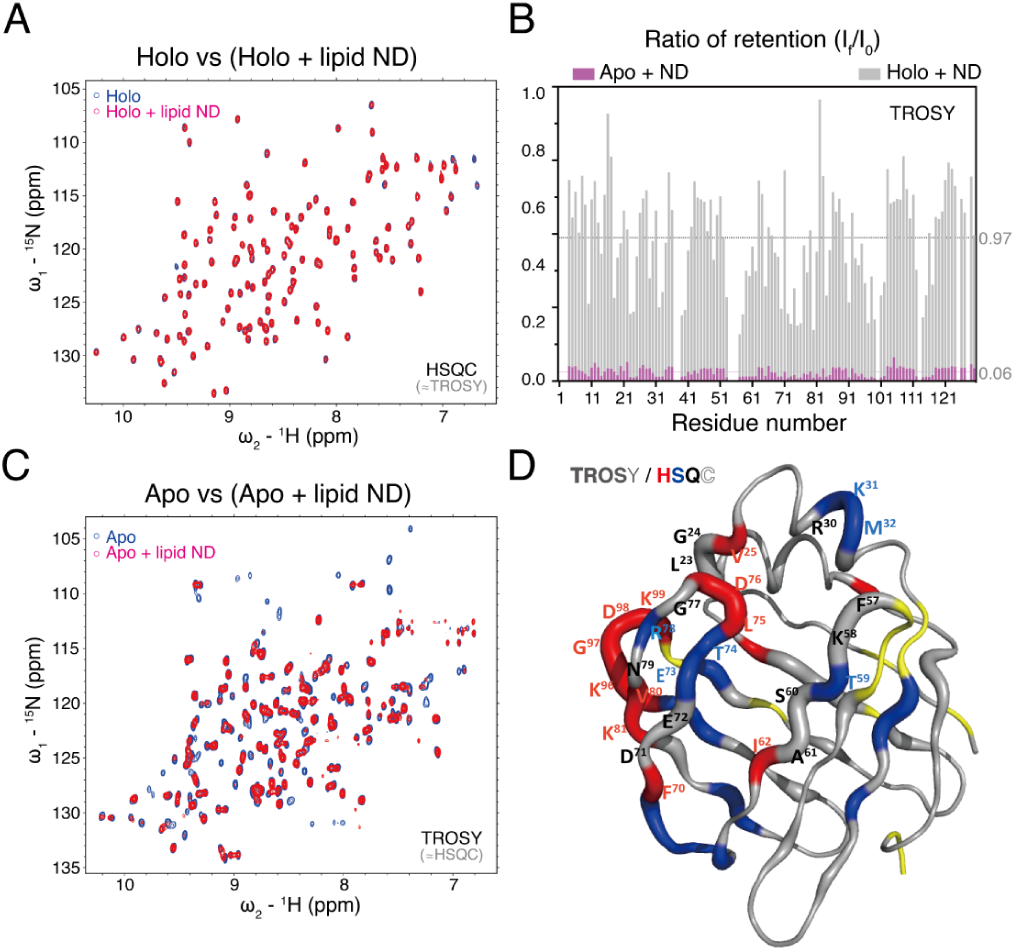
Interaction of GuHCl-purified apo-dFABP (5GKB) with membranes. **(A)** Superimposition of the HSQC spectra of OA-liganded holo-dFABP in the presence (red) or absence (blue) of DOPG/DOPC lipid nanodiscs (ND) at a molar ratio of 1:1. **(B)** The intensity change ratio in the TROSY spectra were obtained by dividing the final intensity (*I*_*f*_, with ND) by the initial inten mapping onto the apo-dFABP structure (5GKB) corresponds the level of signal disappearance (fraction of attenuation in TRnsity (*I*_0_, without ND). The average difference on TROSY spectra between the two states is approximately 10. **(C)** Superimposition of the HSQC and TROSY spectra of apo-dFABP in the presence (red) or absence (blue) of lipid ND at a molar ratio of 1:1. A similar trend was observed for the representative TROSY spectra shown here. **(D)** The thickness of sausage presentatioOSY spectra) after interacting with lipid NDs. Residues in the *α*II, *β*D-F, and *β*G-H loops and in *α*I-II (***see Fig. 3B***) with larger changes are labelled, and the unassigned residues are colored yellow. The thicker segment above the average fraction of attenuation is further divided into three groups according to a serial lipid ND titration (followed using HSQC). Red indicates that the signal decayed immediately, whereas blue indicates that the signal decayed faster than those of the other residues (labelled in black).

The diminishing peaks were clustered near the portal and gap regions and near the ligand entry gates (***Fig. 4D, 5B***). This harmony between membrane and ligand interactions enabled the development of a feasible model in which ligand uptake occurs after the attachment of dFABP to membranes near the *β*G-H loop, and the mechanism involves sliding to the region that encompasses the *α*II and *β*D-F regions. To further decipher the enigmatic membrane interaction, residues with attenuation differences above the average value in the TROSY spectra were selected, and their signal decay rates were analyzed using HSQC lipid ND titration (***Supp. Fig. S5A and B***). After the integrated evaluation, including signal attenuation and decay rate, four clusters with the following characteristics were defined: 1) signals that decayed more slowly than the average rate (e.g., R30, shown in black); 2) signals that decayed faster than the average rate (e.g., T74, shown in blue); 3) signals that immediately disappeared upon the addition of lipid NDs (e.g., K99, shown in red); and 4) signals that remained detectable (e.g., L51, shown in hollow font) (***Supp. Fig. S5B and C***). The red and blue clusters are also shown in ***Fig. 4D***. More polar and charged residues (namely, D76, R78, K81, K96, D98, and K99), which are responsible for primary membrane attachment, were observed, followed by successive sliding to integrate nonpolar and aromatic residues (namely, L23, G24, V25, F57, and A61) in the portal and gap regions. Polar residues can be reasonably inferred to engage the polar heads of biomembranes and nonpolar residues to interact with hydrophobic ligands embedded in lipid bilayers. This interplay of membrane–association regions constructed in space is likely to imitate amphitropic peripheral proteins.

A ligand rescue experiment was performed using OA for further analyses of the cross-peak change, and representative residues participating in the membrane interaction are shown in ***Supp. Fig. S6***. Notably, the signals representing apo-dFABP binding to nanodiscs recovered by rescued ligand OA were more similar for conformer-B’, which did not completely superimpose on the spectra of OA-liganded holo-dFABP, implying that the ligand can be internalized into the escort protein and modulate the dissociation of the holo form from the membrane mimetics. In relation to both ligand binding and membrane interaction, four out of five selected residues shown in ***Fig. 5B*** are polar and charged, and the two residues in squares (R30 and S60) lie at the protein–membrane interface. The unknown ligand map coheres with those of ligands from other similar escort proteins (e.g., heart-type FABP, epidermal-type FABP and liver-type FABP), and the docking complex signifies that the aliphatic tails of ligands around the portal region interfere with the protein–membrane interactions (***Fig. 5C***).

**Figure 5.**
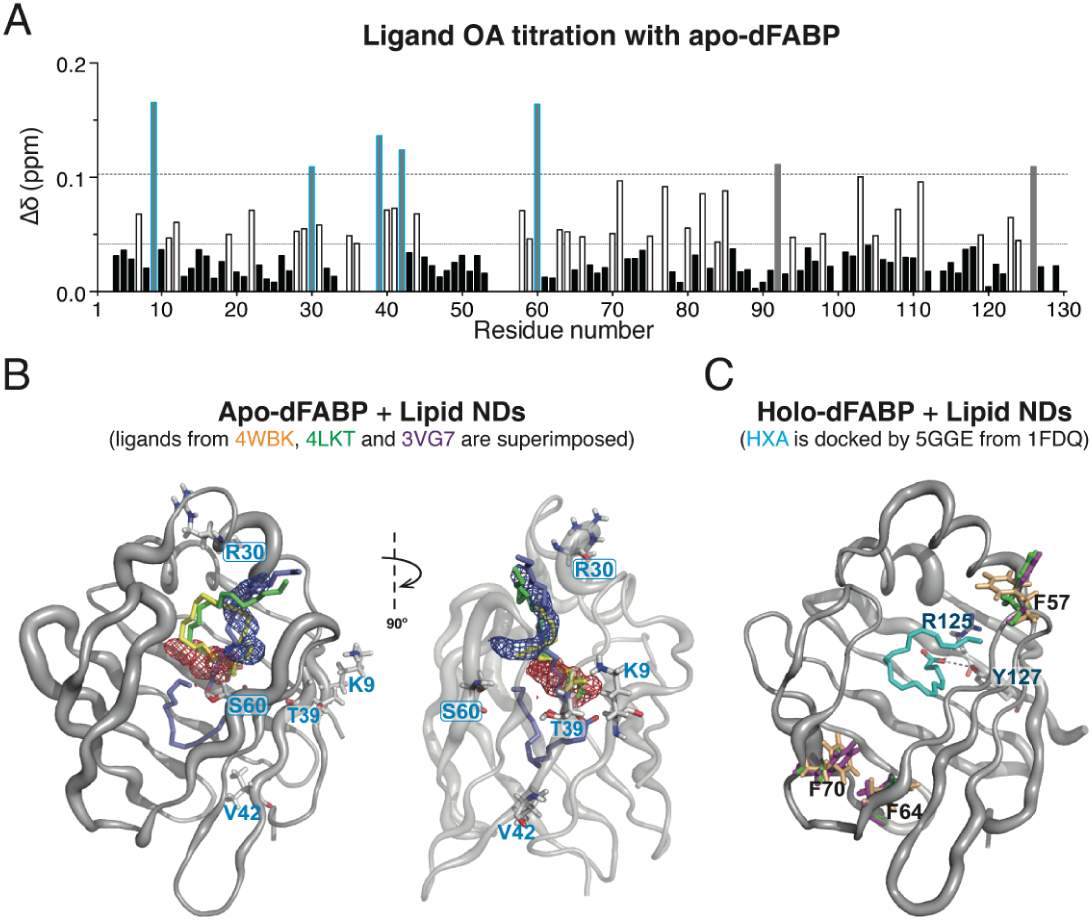
Intersection analysis between ligand binding and membrane interaction. **(A)** The chemical shift perturbations of apo-dFABP were measured by ligand OA titration. Residues with values greater than the total average value (0.04 ppm, the lower, more closely spaced dotted line) were recalculated to obtain an average value similar to that of the upper, more widely spaced dotted line (approximately 0.1 ppm). Five of the seven residues above the highest average CSP value (in blue frames) are found to have higher levels of disappearance (fraction of attenuation in TROSY spectra) during lipid ND interaction. **(B)** B. The effect of lipid NDs on apo-dFABP mapping onto the apo-dFABP structure (5GKB) is shown in sausage representation. The five residues with higher levels of disappearance during lipid ND interaction are shown as sticks and are labelled on the structure; two of these (in squares) are located in the membrane-associated region. The maps of the unknown ligand in liganded-like dFABP (5GGE, red and blue meshes), stearic acid of human heart-type FABP (4WBK, yellow sticks), arachidonic acid of human epidermal-type FABP (4LKT, green sticks) and palmitic acid in human liver-type FABP (3VG7, purple sticks) are shown using the PyMOL structure alignment suit for ligand orientation comparison. **(C)** The effect of lipid NDs on holo-dFABP mapping onto the liganded-like structure (5GGE) is shown in sausage representation. The structure is superimposed with the 5GGE docking ligand using docosahexanoic acid (HXA, cyan sticks) in human B-FABP (1FDQ) and PatchDock, and the two residues interacting with the carboxylate head group of the docked HXA ligand are indicated in CPK with labels. Three aromatic residues of 5GGE, 5GKB and the docking model, which are located in the *β*C-D loop and *β*D-E, are labelled in magenta, green and mustard, respectively.

According to comprehensive data obtained for these residues, when the apo-dFABP first approaches a lipid ND, a rolling motion occurs in the region composing the*β*G-H*β*E-F loops and the *a*I-II turn, which subsequently turns towards the portal and gap regions for ligand accommodation (***Fig. 6***). In particular, the intensity of conformer-B’ remained almost unchanged during lipid ND titration, whereas that of conformer-A of apo-dFABP continued to diminish (***Supp. Fig. S6B and S7C***). Together with the consistent results showing a barely changed intensity in conformer-B’ between apo-dFABP and holo-dFABP during lipid ND interaction, these findings strongly indicate that the ligands escorted by dFABP would favour unidirectional transportation (***Fig. 4A and B***).

**Figure 6.**
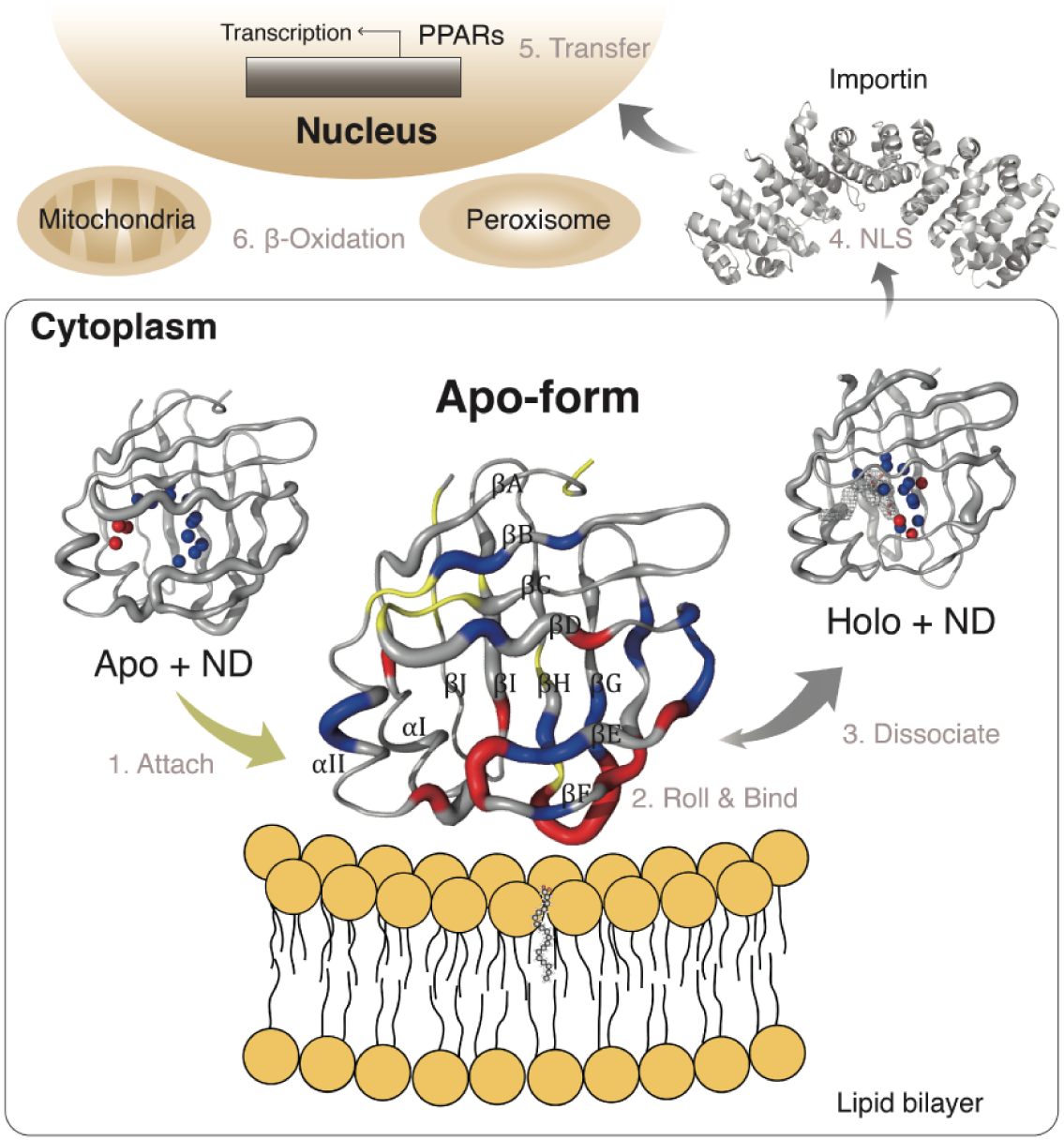
Proposed model for ligand-regulated dFABP transport. The mechanistic trafficking model of fatty acid (FA) accompanied by dFABP is shown in detail. A putative ligand transfer model was proposed in conjunction with our study (as shown in the square) as follows: 1) The apo form of the escort protein FABP attaches to the lipid bilayer. 2) Apo-FABP slides to accommodate complementary LCFAs. 3) The ligand-bound form, holo-FABP, dissociates from the lipid bilayer. 4) Importin *α*(PDBID: 1EJL) recognizes the spatial arrangement of the NLS in the holo form of FABPs (***Amber-Vitos et al., 2015***). 5) Holo-FABP transfers the ligand to PPARs (***Armstrong et al., 2014***). 6) PPARs trigger gene transcription and produce proteins for lipid metabolism (e.g., *β*-oxidation) in the mitochondria or through other signaling pathways. The fraction of attenuation is illustrated in sausage representation for apo- and holo-dFABP (***Fig. 4D, Supp. Table S14***) (5GKB and 5GGE, respectively), and the PyMOL-defined cavity water molecules and incomplete ligand-like maps are shown in spheres and meshes, respectively.

## Discussion

Intracellular cargo transport is a complex and highly dynamic process. By analyzing dFABP with enriched conformational homogeneity, we present a mechanistic model for the transport directionality of lipophilic cargo molecules in representative escort proteins, namely, FABPs. FABPs comprise ten *β* strands, which are organized orthogonally as a clamp appended to a short helix-turn-helix motif as the portal cap region near N terminus (33). Previous studies have reported that the minor open state of the portal region in apo forms of FABPs was responsible for ligand entry (17, 34), while the slow protein conformational exchange was suggested to be irrelevant to the ligand association (18). A fundamental problem with escort protein-dependent ligand transport, such as the transport of LCFAs by FABPs, is whether the directionality is reversible or irreversible. Initial analysis of the ligand–FABP complex structures in the Protein Data Bank did not reach any conclusion regarding the transport mechanism (35), and the hypothesis that the inward state of the conserved Phe residue in the *β*C-D loop of FABPs corresponds to the liganded closed form and the outward state to the unliganded open form has not been universally consistent (e.g., 5BVQ, 4I3B). In our view, the potential interactions among several key factors, including the escort protein, the target ligand, and the ligand donor, all need to be considered concurrently for a successful interpretation of the dynamic cargo transport mechanism.

In this article, we provide several lines of compelling evidence, including biophysical analyses and structural comparisons, to support our speculation that the lipophilic ligand facilitates protein–ligand complex dissociation from ligand donor membranes. The level of trace contaminants in the urea-treated sample was unproblematic in our conventional ligand-protein interaction analyses (***Supp. Fig. S4B and C, Supp. Fig. S8***). According to the CD studies shown in ***Fig. 1*** and ***Supp. Fig. S2***, however, some biophysical properties such as thermal stability might yield inappropriate interpretations (36, 37). The decreased Tm of the urea-treated sample in CD experiments might have resulted from an association between the protein and nonprotein membrane-like debris (***Supp. Fig. S3***). The differences between the apo and holo forms of urea-treated dFABP that are seen in far-UV CD spectra may result from the presence of associated impurity because both forms had secondary structures with similar thermal stability in the GuHCl-purified sample (***Supp. Fig. S2A***). Further investigation of the membrane interactions confirmed that the holo form had far fewer interactions with membrane mimetics (***Fig. 4A and B***) and that the apo form dissociated from membranes by an additional ligand binding, as seen in the rescue study (***Supp. Fig. S6C***). Therefore, the ability of OA to cause the urea-treated sample to dissociate from membrane-like debris and adopt the free OA-liganded form would explain the higher secondary thermal stability of the OA-liganded urea-treated dFABP (***Supp. Fig. S2A***). Although the analysis of protein thermal stability based on precipitation using far-UV CD spectra was not applicable (***Supp. Fig. S2A and D***), the proteins unfolded cooperatively and globally, as seen in intrinsic fluorescence experiments (***Supp. Fig. S2C***). The changes in Tm (as determined using intrinsic fluorescence spectra) between the apo and holo forms of urea-treated and GuHCl-purified dFABP were approximately 10 °C and 5 °C, respectively, and both holo forms had identical melting curves, suggesting that the liganded forms of dFABP were consistent (***Supp. Fig. S2C***). Moreover, the difference in stability between the apo and holo forms was confirmed by a chemical denaturation experiment, which showed that the rigidity assisted by the ligand was similar to the thermal stability (***Supp. Fig. S4A***). Altogether, the intrinsic fluorescence experiments on dFABP demonstrated that the 1ndings regarding the difference in thermal unfolding between the apo and holo forms were consistent with those of previous CD studies (37, 38), even if the Tm values from the CD spectra were not readily interpreted.

We discovered through both microscopic and macroscopic analyses that the general heterogeneity of FABPs, including unknown ligands and membrane-like debris, may be a reason for inconsistency in the Phe residue orientation in the *β*C-D loop (e.g., penguin apo-AFABP, 5BVQ). Based on the crystallization results, the urea-treated liganded-like dFABP exhibited a tertiary structure similar to that of the GuHCl-purified apo-dFABP, except that no ligand-like electron density was calculated inside its binding cavity (***Fig. 3***). Therefore, the difference in temperature factors around the portal and gap regions may be attributed to conformational heterogeneity (***Supp. Fig. S10***). On the basis of a further comparison of the ordered waters in the ligand binding cavity of liganded-like and apo-dFABPs, we conjectured that the residues around ligand entry sites likely have higher temperature factors in the apo form of urea-treated dFABP because of the possible lack of an ordered-water hydrogen networks resulting from the flexible hydrophobic tail of the unknown ligand (***Fig. 3A***). Although the HSQC spectrum of urea-treated liganded-like dFABP exhibited minor perturbation upon ligand binding (***Supp. Fig. S9A***), a high proportion of the holo form left inadequate space for further membrane interaction. The spectra of urea-treated liganded-like dFABP and OA-dFABP, especially in the region from *β*D to E, are indistinguishable from ligand entry to membrane interaction (***Fig. 2D and 4D***).

Consequently, GuHCl-purified apo-dFABP was used to study the relationship between proteins and membranes in the presence and absence of ligands. The broadened apo-dFABP signals resulting from membrane interaction were superimposed on the unliganded 5GKB structure, and the residues were grouped by decay features corresponding to the order of membrane attachment (***Fig. 4D, Supp. Fig. S5***). By carefully dissecting the clusters in terms of ligand binding and membrane interaction, a harmonious coupling was found to explain the interactions among lipophilic ligands, escort proteins, and bilayer membranes as follows: 1) based on the crystallographic temperature factors, we assumed that *β*E-F controls ligand entry; and 2) membrane attachment was described as being initiated by attaching to the *β*G-H loop and rolling towards the region involving the *α*II and *β*D-E because the signals for the *β*G-H and *β*E-F loops decayed faster than those for the portal and gap regions (*α*II/*β*D-E) (***Fig. 6***). The ligand is assumed to participate in the interface between the protein and the membrane, and liganded dFABP tended to dissociate from the membranes after binding to the ligand, as reported by ligand OA rescue analysis (***Supp. Fig. S5***).

Numerous studies have suggested that positively charged residues in the helix region control membrane interactions (39-41). More recent research has demonstrated that these residues are part of a spatially arranged nuclear leading sequence (42-44). Although residues in *α*II, such as R30, were disturbed by ligand and membrane titration (***Supp. Fig. S6A and B***), they were neither the main cause of nor independent contributors to the membrane attachment. In addition to allowing further study of membrane interaction, the enriched conformational homogeneity of dFABP provides a promising means of distinguishing membrane interactions from other interactions and demonstrates that the conformer-A signals of residues that participate in membrane interaction tend to disappear when the interaction occurs (***Supp. Fig. S6B, S7C***). Residues that do not strongly contribute to membrane interaction and those facing the ligand binding cavity (e.g., Y127) do not fully lose intensity or show no chemical shift perturbation after membrane titration (***Supp. Fig. S7D***), in contrast to how ligand titration transforms conformer-A into the OA-liganded holo form (***Supp. Fig. S7B***).

According to our findings, the greatly decreased membrane affinity of liganded dFABP determines ligand transport directionality. Not only does the orientation of ligand binding inside the protein affect repulsion in the protein–membrane interaction (***Fig. 7B and C***), but also the ligand donor membranes may induce the open state of the escort protein to enable ligand uptake (34). Considering that lipophilic LCFAs are physiologically embedded in membranes and that high affinity for FABPs is observed in vitro, the combination of these two scenarios may further explain why not all ligands with similar binding affinities can trigger FABP nuclear translocation. First, ligands that can be uploaded complementarily into the escort protein may repel either the *α*II of FABPs (45) or the charged interface of FABP–membrane due to their aliphatic tails and thereby force FABPs to dissociate from the membrane (***Fig. 5***). In contrast, ligands that do not fold properly into the binding cavity of FABPs would leave a tail below the membrane surface inside the hydrophobic environment, thus preventing the protein from dissociating. Moreover, the spatial distribution of charged residues in *β*C-F and the conserved Phe residue in the *β*C-D loop seems to facilitate the attachment of FABPs to membranes. These findings might be crucial, necessitating the re-evaluation of previous studies on the interactions among FABPs, complementary ligands and membranes based on molecular dynamic simulations, which have mostly focused on explaining how LCFAs enter FABPs or how holo-FABPs transport ligands to membranes (46-48). The relationship between liganded FABPs and membranes provides an alternative interpretation: only through a specific orientation of the ligand in the protein can the aliphatic tail modulate protein–membrane dissociation and act as activator ligands to control protein–ligand complex release for nucleocytoplasmic translocation (44). The use of blockers targeting holo-FABPs instead of competitors for apo-FABPs may be able to interrupt the pathways for entering the nucleus or for interacting with effectors. Through illuminating the unidirectional transport by dFABP, this study advances research on FABPs, particularly on their relationships with complementary LCFAs and membranes, which will enable further investigation of their roles in the nucleus. The mechanism generalizes a model, obtained by clarifying the working patterns of FABPs, in which the complementary lipophilic ligand is the decisive factor in regulating its own transport directionality in membrane-associated escort proteins.

## Materials and Methods

### Cloning and expression of dFABP

The second spliced isoform (CG6783-B) of the dFABP locus (CG6783) in the *Drosophila* genome was found to share the highest homology to mammalian brain-type FABP, and its 130-residue peptide sequence was expressed and purified for subsequent structural studies (49). In brief, the open reading frame coding for dFABP isoform B was chemically synthesized with E. coli-optimized codons (Eurofins Scientific, Luxembourg), and inserted into the pET23a(+) expression vector (Merck, Darmstadt, Germany) at the NdeI/XhoI sites. The dFABP-pET23a plasmid was subsequently transformed into E. coli BL21-Gold (DE3) competent cells (Thermo Fisher Scientific, MA, USA) for recombinant protein expression. Individual colonies were selected, and an aliquot of host cells harboring the correct insert was inoculated into 20 mL of Luria broth (LB) containing 100 *µ*g/mL ampicillin. The broth was incubated at 37 °C with 180 rpm agitation for 12 h as a condensed starting culture. The 20-mL cell culture was either poured into 800 mL of LB or precipitated and then resuspended in 1 L of uniformly isotopes-labelled M9 medium, which consisted of 1 g of ^13^C-glucose/1 g ^15^N-NH_4_Cl (Cambridge Isotopes Laboratories, MA, USA) with 20 *µ*g/mL ampicillin. Subsequently, dFABP expression was induced with 0.2 mM isopropyl *β*-D-1-thio-galactopyranoside (IPTG) when the optical density at 600 nm reached 0.6, and the cultures were incubated for an additional 6 h at 37 °C.

### Purification of dFABP with higher conformational homogeneity

***Supp. Fig. S1*** represents the flowchart for dFABP purification, by our improved new strategy with the combination of 8 M urea and the partially unfolding reagent 2 M GuHCl, depending on the denaturation analysis (***Supp. Fig. S4A***). Brie2y, a batch of 800 mL LB broth or 1 L isotopes-labelled of the protein expression culture was harvested by 12,000 × *g* centrifugation at 4 °C for 20 min. Cell pellets were resuspended collectively in 30 mL of lysis buffer (20 mM Tris pH 8.0, 2 mM EDTA, 1 mM *β*-ME, 0.05% NaN_3_) and frozen at -20 °C before further purification. The thawed cells were lysed by sonication and centrifuged at 38,500 × *g* for 15 min at 4 °C. The pellet was solubilized in 5 mL of 8 M urea at 4 °C for 3 h and then centrifuged again at 38,500 × *g* at 4 °C for 15 min to remove insoluble debris. The supernatant was dialyzed against 1 L of the sample buffer (20 mM Tris and 1 mM *β*-ME, pH 8.0) using 3-kDa molecular weight cutoff dialysis tubing (Orange Scientific, Braine-l’Alleud, Belgium) and stirred at 4 °C for 6 h. The precipitates were removed by centrifugation at 38,500 × *g* at 4 °C for 15 min, and the supernatant was mixed with an equal volume of buffer B (1 M (NH_4_)_2_SO_4_ in 20 mM Tris, pH 8.0) and concentrated to 10 mL for hydrophobic interaction chromatography (5-mL HiTrap phenyl HP column, GE Healthcare, IL, USA). Subsequently, a 2-mL elution of hydrophilic flow-through was incubated in the absence (denoted as urea-treated dFABP) or presence (denoted as GuHCl-purified dFABP) of 2 M GuHCl preincubated at room temperature for 30 min for the next size-exclusion chromatography. The urea-treated dFABP was obtained and validated by twice injection of S-200 size-exclusion chromatography (HiPrep 16/60 Sephacry S-200 HR, GE Healthcare), and the GuHCl-purified dFABP underwent two rounds of S-200 size-exclusion chromatography, with final validation by S-75 size-exclusion chromatography (Superdex 75 10/300 GL, GE Healthcare). All chromatography samples had a volume of 5 mL after the removal of precipitates by 0.22-*µ*m membrane filters. The crude quality of the desired dFABP protein after each step was assessed using SDS-PAGE. The polished samples at the final purification step were further determined through ESI UPLC Q-TOF mass spectrometry (Impact HD, Bruker, MA, USA) and the yield of 1 L of culture was approximately 15 mg, as measured by Bradford and BCA assays. The final dFABP samples were stored in 500 *µ*M in the sample buffer at 4 °C until use in 6 months.

### Spectroscopic characterization of dFABP

A 25-*µ*M sample was prepared in sample buffer using 0.1-cm quartz cuvette for an Aviv Model 202 CD spectrometer (Aviv Biomedical, NJ, USA) and a 2-*µ*M sample was used for intrinsic fluorescence measurement (F-7000, HITACHI, Tokyo, Japan). The mean residue ellipticity (MRE) was calculated based on the initial sample concentration in temperature-dependent far-UV CD experiments. As a result of the decrease in soluble concentration during the heating process in the urea-treated sample, the CD melting curves from 25 °C to 95 °C measured at a wavelength of 215 nm were expressed in millidegrees instead of MRE. Holo-dFABP was combined with OA (100 mM stock in 250 mM KOH, Avanti Polar Lipids, AL, USA) to achieve a molar ratio of 1:2 in the CD and fluorescence spectroscopic experiments. The excitation wavelength of intrinsic fluorescence was set at 275 nm because there were four tyrosines instead of tryptophan in the dFABP sequence and detected at an emission wavelength of 303 nm. Each temperature of the stepwise melting experiments was equilibrated for 10 s before manual detection. The CD melting curve was automatically set at increments of 2 °C from 24 °C to 96 °C and equilibrated for 1 s with 0.5-s data averaging. The assay results were fitted using the Sigmoidal model in GraphPad Prism 6 software (GraphPad Software, Inc. La Jolla, CA, USA).

### Crystallographic characterization of dFABP

The dFABP samples were concentrated to approximately 8 mg/mL in 20 mM Tris buffer (pH 8.0) for crystallization. Crystals were screened either manually or by using the automatic protein crystallization robot Oryx8 (Douglas Instruments, Hungerford, UK) with Wizard crystallization screens (Rigaku, WA, USA). The urea-treated dFABP (denoted as 5GGE) was crystallized through vapour diffusion in hanging-drop format, and a 1-*µ*L protein sample was mixed with a 1-*µ*L drop of reservoir solution against 200 *µ*L of reservoir solution in 24-well Linbro plates (Flow Laboratories, VA, USA). The GuHCl-purified dFABP (denoted as 5GKB) was crystallized through vapour diffusion in sitting-drop format. The microbatch screening system was applied under the crystallization conditions of Emerald Wizard II-F7 (0.1 M phosphate citrate, 1.6 M NaH_2_PO_4_/0.4 M K_2_HPO_4_, final pH 5.2), and 0.3 *µ*L of GuHCl-purified dFABP was mixed with 0.3 *µ*L of reservoir solution against 60 *µ*L of crystallization solution in 96-well crystallization plates. To acquire X-ray diffraction data, crystals were grown for 2 months at 20 °C, and data acquisition was performed at the National Synchrotron Radiation Research Center, Taiwan. The data were processed and scaled using HKL2000 software, and the molecular replacement program MOLREP in CCP4 was applied for phase determination (50). The initial models for 5GGE and 5GKB were 1FTP and 5GGE, respectively. Further model building, the incorporation of the ligand citrate in 5GGE, and both structural refinements were performed using the PHENIX suite (51). The structure of liganded-like dFABP (5GGE) was subjected to the docking program PatchDock for ligand docking (docosahexanoic acid, HXA of 1FDQ).

### NMR HSQC spectroscopy analysis

Samples for NMR were buffer exchanged into 50 mM phosphate buffer (pH 6.8) with the addition of 10% D_2_O and trace 4,4-dimethyl-4-silapentane-1-sulfonate as a calibration standard for spectra collected at 298K. An Avance-600 MHz spectrometer was used for the OA titration of HSQC spectra, and an Avance-850 MHz spectrometer (Bruker, Germany) equipped with a cryoprobe was used to acquire the spectra for nanodisc interactions. The holo form of the OA-dFABP sample was prepared at a 1:3 molar ratio of protein to OA, and the 3D spectra (HNCA, HNCOCA, HNCACB, CBCACONH, HNCO and HNCACO) for sequential assignments were acquired on the Avance-850 MHz instrument. Identical 3D NMR experiments for assignments of apo-dFABP were recorded for back-calculation from the OA-dFABP assignments (Supp. Table S11, S12). The data were processed using NMRPipe (52) and autoassigned by MANI PINE server v.2.0 (53), with manual inspection through NMR assignment and Sparky integration software (Goddard TD & Kneller DG, 2008, SPARKY 3. University of California, San Francisco, CA, USA). Chemical shift perturbations (CSPs) from the initial conformer-A to the final holo form for dFABP were analyzed by weighting the averaged ^1^H and ^15^N chemical shift changes with the equation Δ*δ* = [(Δ*δ*_1*HN*_)^2^ + (0.17Δ*δ*_15*N*_)^2^]^1/2^.

### Membrane interaction of dFABP with lipid bilayer nanodiscs

The scaffold protein MSP1D1 was expressed and purified for the self-assembly of lipid bilayer nanodiscs with DOPG/DOPC (1:1) as a membrane mimetic (54). GuHCl-purified dFABP was prepared at 100 *µ*M and 70 *µ*M for lipid ND titration, and HSQC and TROSY spectra were obtained, respectively. Holo-dFABP was mixed with 2.5-fold OA to ensure ligand saturation before interaction with lipid NDs. A final 1:1 ratio of protein to lipid NDs was used for the TROSY and HSQC spectra. The settings for the TROSY spectra were based on the pulse sequence method described previously (55) in order to optimize the power of water suppression by gradient-tailored excitation. The data calculation for the fraction of attenuation was weighed by dividing the initial intensity by a factor 10, depending on the diminishing effect in TROSY spectra, to enlarge the difference in the broadened signals after the addition of lipid NDs to apo-dFABP.

## Acknowledgements

We thank the Biological Crystallography Faculty of National Synchrotron Radiation Research Center (Taipei, Taiwan) for technical supports and assistance in the X-ray and data collection. We are grateful to Dr. Tsyr-Yan Dharma Yu and Mr. Julien Massiot of the Institute of Atomic and Molecular Sciences at Academia Sinica (Taipei, Taiwan) for constructive guidance on TROSY experiments and kind preparation of lipid nanodiscs. We appreciate Dr. Yet-Ran Chen of the Agricultural Biotechnology Research Center at Academia Sinica (Taipei, Taiwan) for valuable comments on mass spectra and lipidome experiments. We are indebted to Drs. Lee-Wei Yang and Shih-Che Sue of the Institute of Bioinformatics and Structural Biology at National Tsing Hua University (Hsinchu, Taiwan), and Dr. Shang-Te Danny Hsu of the Institute of Biological Chemistry at Academia Sinica (Taipei, Taiwan) for their insightful remarks and helpful suggestions on this project. We are also thankful to Dr. Yi-Zong Lee and Mr. Liang-Yuan Chiu of the Institute of Bioinformatics and Structural Biology at National Tsing Hua University (Hsinchu, Taiwan) for their kind assistance with 3D NMR spectra. This research was supported by grants from Ministry of Science and Technology (MOST106-2319-B-400-001; MOST105-2311-B-007-002 & MOST103-2325-B-400-002) and National Health Research Institute (CA-103-PP-13; CA-104-PP-13 & CA-105-PP-13), Taiwan.

## Additional information

### Author contributions

Y.Y.C. designed the study, executed all experimental work, data interpretation and drafted the manuscript. Y.F.H. collaborated in improving the protein purification processes. H.H.L. contributed to cloning and crystal collection of urea-treated wild-type dFABP. W.S.C. and P.C.L. contributed to project supervision and assisted manuscript preparation.

---

**Supplementary Fig. S1.**
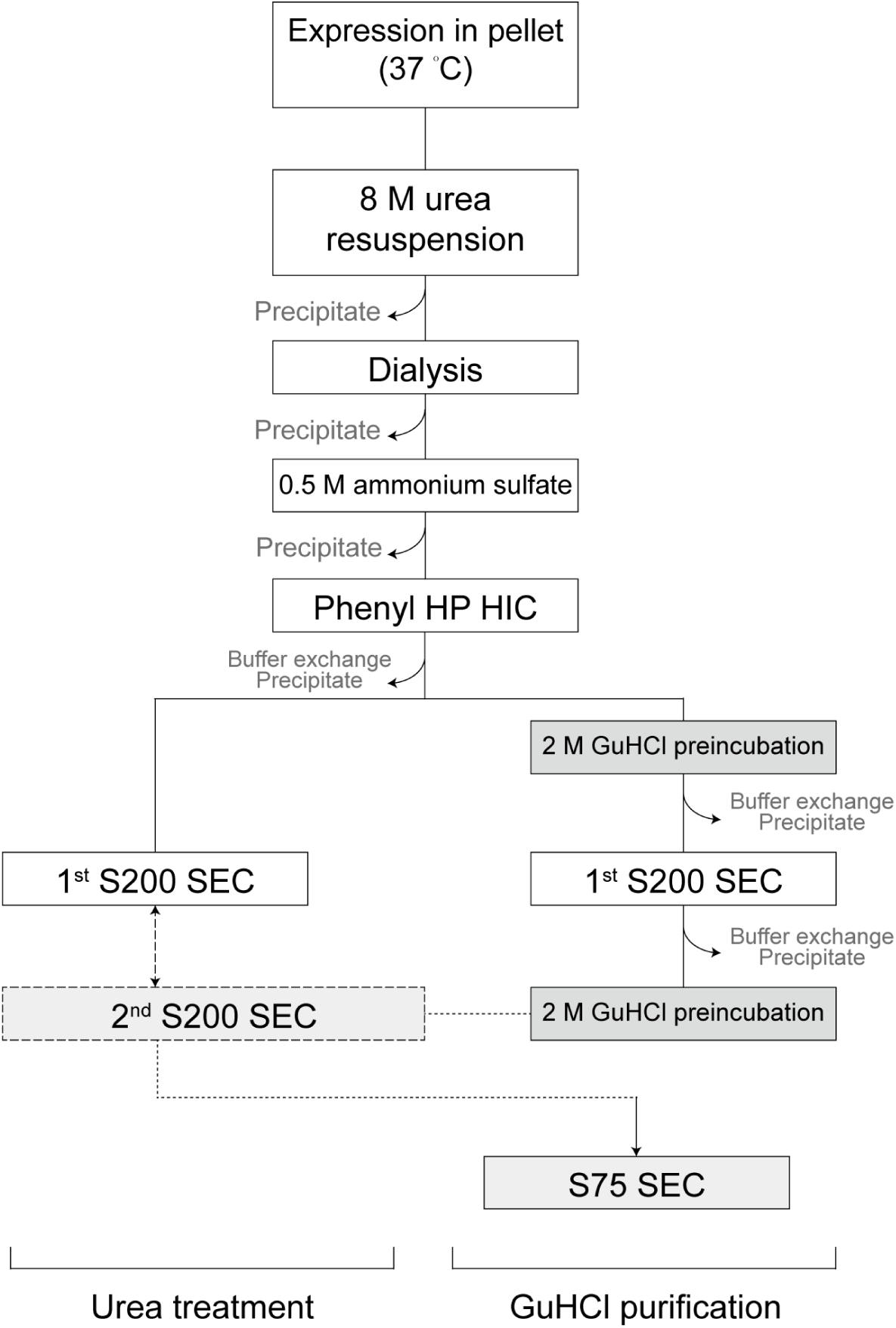
Scheme of the purification procedures for urea-treated and GuHCI-purified dFABP.

**Supplementary Fig. S2.**
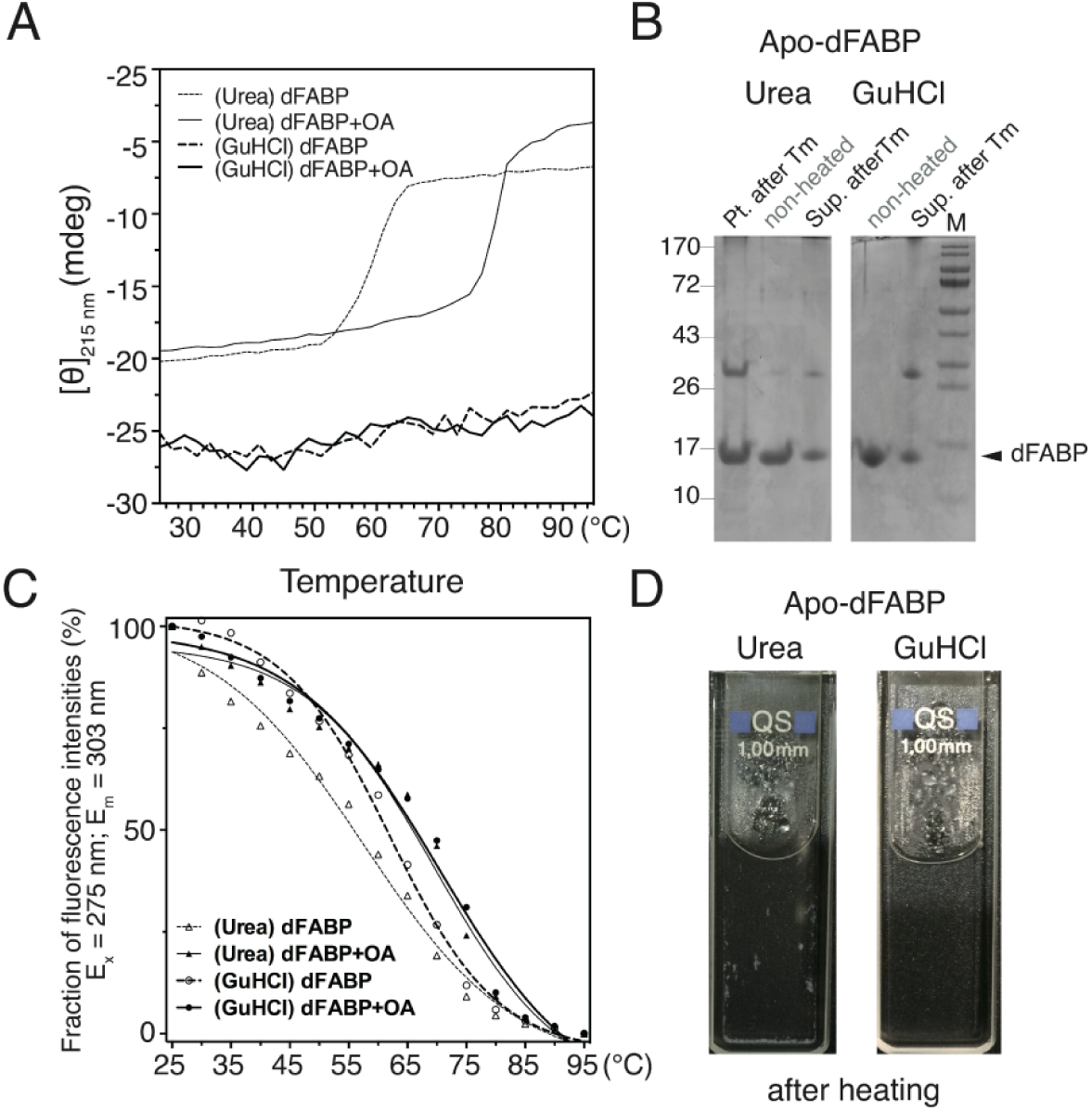
Comparison of thermal stability and sample solution after heating. **(A)** CD thermal melting curves of urea-treated and GuHCl-purified samples were compared in the presence or absence of ligand OA. No apparent secondary conformational change was detected at 215 nm for the GuHCl-purified dFABP with or without the ligand OA. **(B)** SDS-PAGE analyses of the urea-treated and GuHCl-purified samples before and after heating in CD Tm experiments. The urea-treated sample showed solid precipitates after heating, whereas the GuHCI-purified sample showed a clear solution, with both supernatants displaying bands at higher molecular weights. **(C)** The unfolding curves derived from the intrinsic fluorescence spectra of urea-treated and GuHCI-purified dFABP in the presence or absence of the ligand OA were measured to determine the tertiary thermal stability. **(D)** The sample solution of GuHCl-purified apo-dFABP remained transparent after heating, whereas that of urea-treated apo-dFABP formed white solid precipitates.

**Supplementary Fig. S3.**
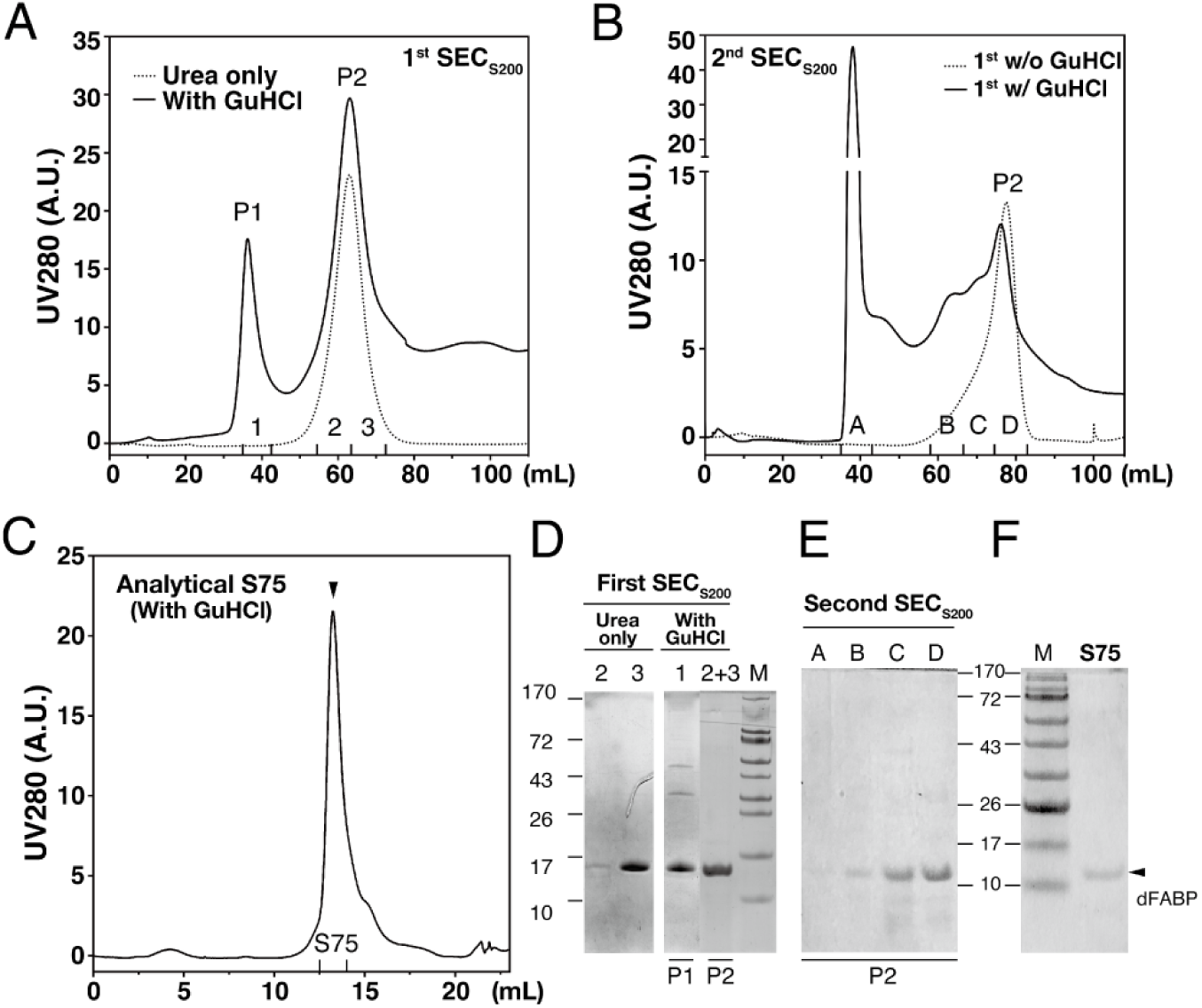
Comparison of the purification profiles of urea-treated and GuHCI-purified dFABP. **(A)** The first round of S200 size-exclusion chromatography (SEC) for HIC eluted samples is presented with (solid line) and without (dotted line) the 2 M GuHCl treatment. With the treatment, two groups of components were separated and denoted eluent P1 (1) and eluent P2 (2 and 3). **(B)** Eluent P2 of the GuHCl-purified dFABP from the first S200 SEC in the presence (solid-line) or absence (dotted line) of repeated 2 M GuHCl treatment is compared for the second round of S200 SEC. This step resulted in the separation of four components, denoted as eluents A, B, C and D. **(C)** Eluent D was finally confirmed by analytical S75 chromatography to be a single elution peak. The sample was preincubated with 2 M GuHCl before injection. **(D)** Eluents of the first round of S200 SEC from urea-treated and GuHCl-purified samples were analyzed using SDS-PAGE and denoted as 1 (P1), 2 and 3 (P2). **(E)** SDS-PAGE analysis of the eluents in the second round of S200 SEC involving GuHCI-purified samples. Four isolated eluents (denoted A, B, C and D) were separated by repeated preincubation with 2 M GuHCl before the injection. **(F)** SDS-PAGE determination of the final fraction of S75 chromatography from the GuHCl-purified sample. The arrow indicates protein bands corresponding to the representative escort protein dFABP.

**Supplementary Fig. S4.**
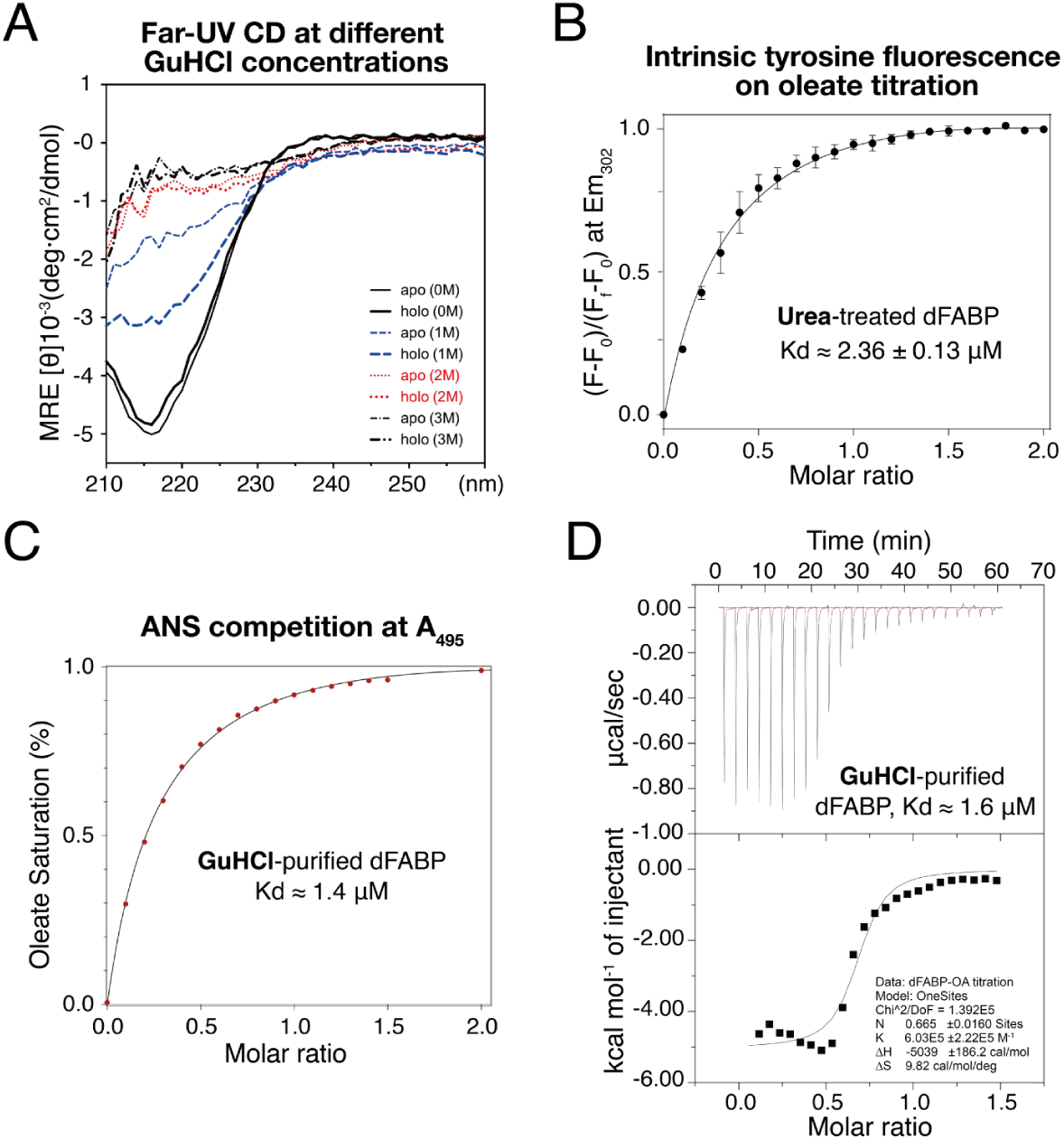
Biochemical properties of dFABP. **(A)** Urea-treated dFABP was titrated with GuHCl (0 M to 3 M) and monitored by far-UV CD. Both apo- and holo-dFABP were unfolded in 2 M GuHCl, and this concentration was used in the GuHCI purification. **(B)** OA binding assay based on the intrinsic fluorescence of urea-treated dFABP (5GGE): 6 *µ*M protein was mixed with OA at a final molar ratio of 1:2. Data were collected using an excitation wavelength of 280 nm and an emission wavelength of 302 nm. The Kd was calculated to be approximately 2.4 *µ*M. **(C)** ANS competition assay; 1 *µ*M GuHCl-purified dFABP (5GKB) presaturated with 5 *µ*M ANS was mixed with protein in competition with the ligand OA at a final molar ratio of 1:2, and ANS fluorescence (detected at 495 nm) was measured to enable the calculation of Kd calculation (approximately 1.4 *µ*M). **(D)** GuHCl-purified dFABP at a concentration of 170 *µ*M was subjected to iTC200 and titrated with OA at 30°C. The derived Kd is approximately 1.6 *µ*M.

**Supplementary Fig. S5.**
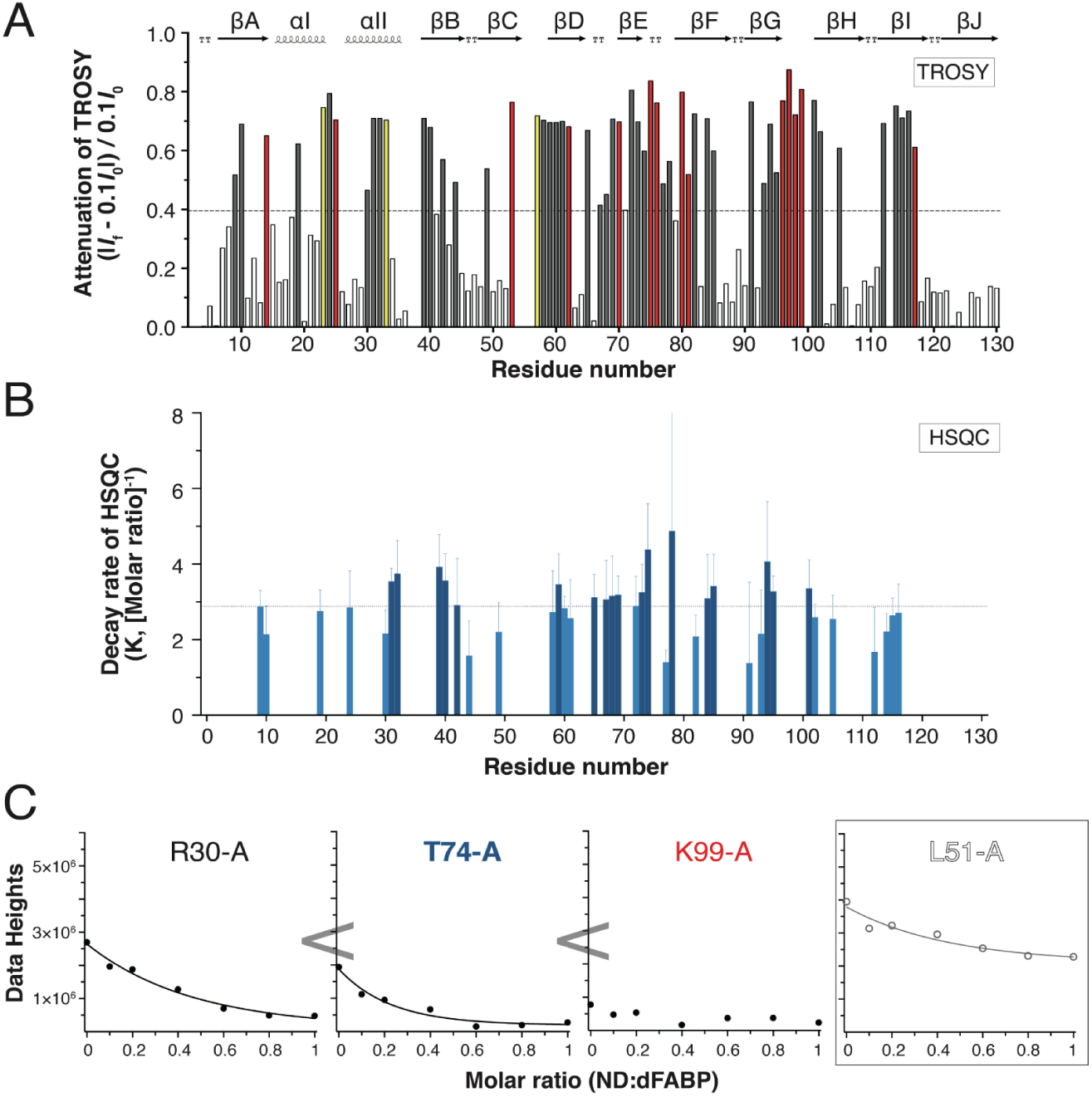
Membrane interaction pattern of apo-dFABP. **(A)** The fraction of attenuation was derived from TROSY spectra of 5GKB in the presence or absence of lipid ND (***Fig. 4C***). The average change is indicated by the dotted line. Residues labelled in red are decayed immediately; others above the average value were analyzed further by lipid ND titration using a series of HSQC spectra. Residues shown in yellow are not assigned. **(B)** Residues with attenuation fractions above the average in the TROSY spectra were selected, and their decay rates were analyzed using lipid ND HSQC titration. The signals of selected residues were fit to the one-phase decay model in GraphPad, and those above the average K (2.88 M^-1^) were defined as faster decaying residues. **(C)** The residues were classified according to a combination of the attenuation fraction and signal decay rate, and representative residues appended with A indicate conformer-A. Residues with consistent intensities bracketed as L51-A are depicted in a thin sausage representation, as shown in ***Fig. 4D***. A single-exponential decay function was applied to all TROSY-selected residues, and three representative residues, namely, R30, T74 and K99, are plotted and colored according to their decay patterns.

**Supplementary Fig. S6.**
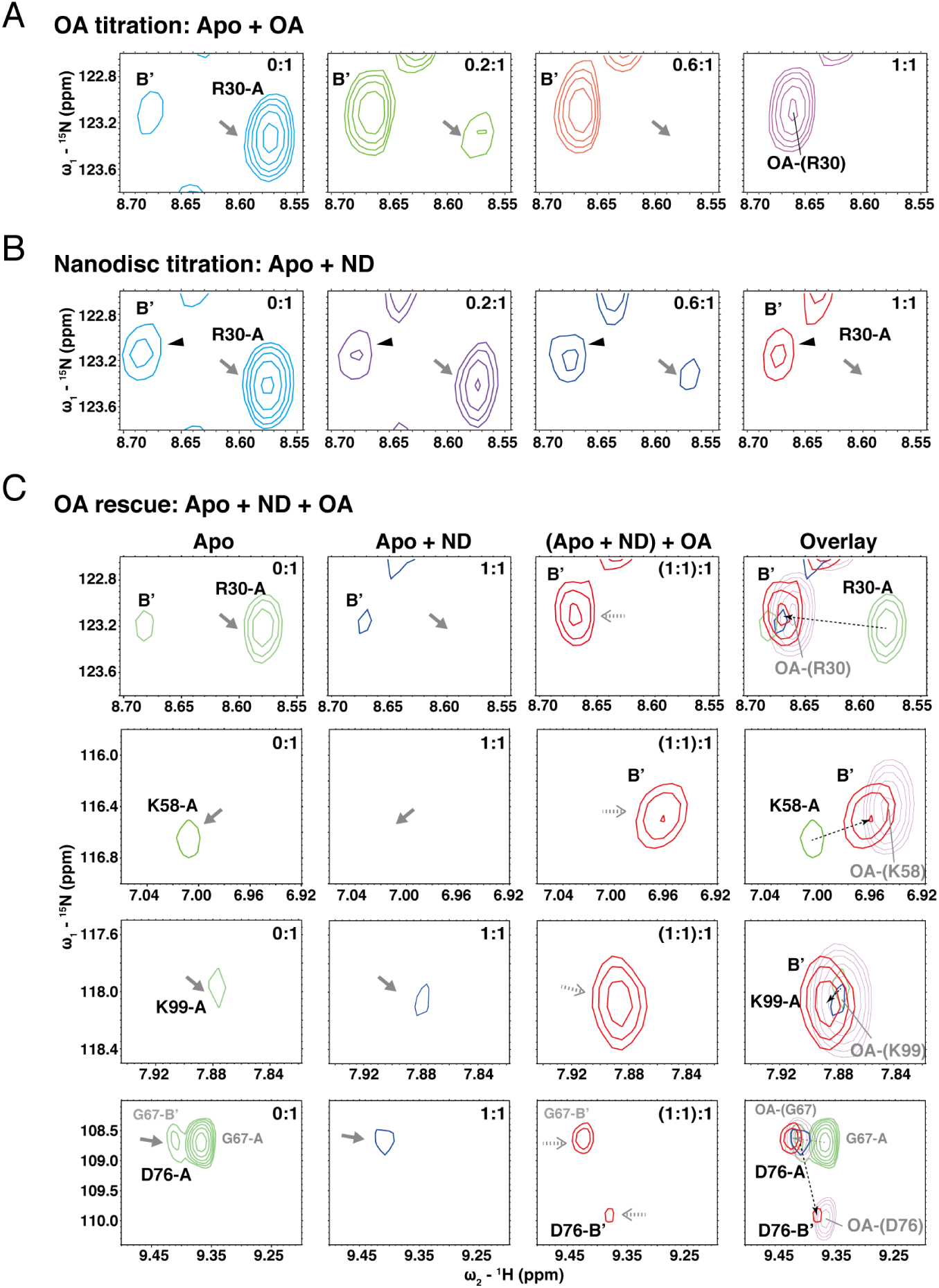
Comparisons of ligand binding and membrane interaction as analyzed in the signal recovery study. **(A)** Four representative ratios in the ligand titration illustrated by R30. The gray arrow indicates the disappearance of conformer-A upon ligand titration, and in the final spectrum, the OA-liganded cross-peak in labelled with the representative residue in brackets. **(B)** Four representative ratios in the lipid ND titration are illustrated by R30. The signal for conformer-B’ of R30 in apo-dFABP changes very little (black arrow), whereas the signal for conformer-A continues to broaden (gray arrow). **(C)** Four representative residues with higher levels of disappearance, as seen in ***Fig. 4D***, in an OA rescue experiment. The signals of conformer-A (gray arrow) of apo-dFABP that disappeared after interaction with membrane mimetics are recovered as conformer-B’ (gray dotted arrow) after the addition of ligand OA. The final comparison panel plots the preceding spectra together with the OA-liganded spectrum (light magenta), and a dotted arrow links the initial (green) and the rescued (red) signals.

**Supplementary Fig. S7.**
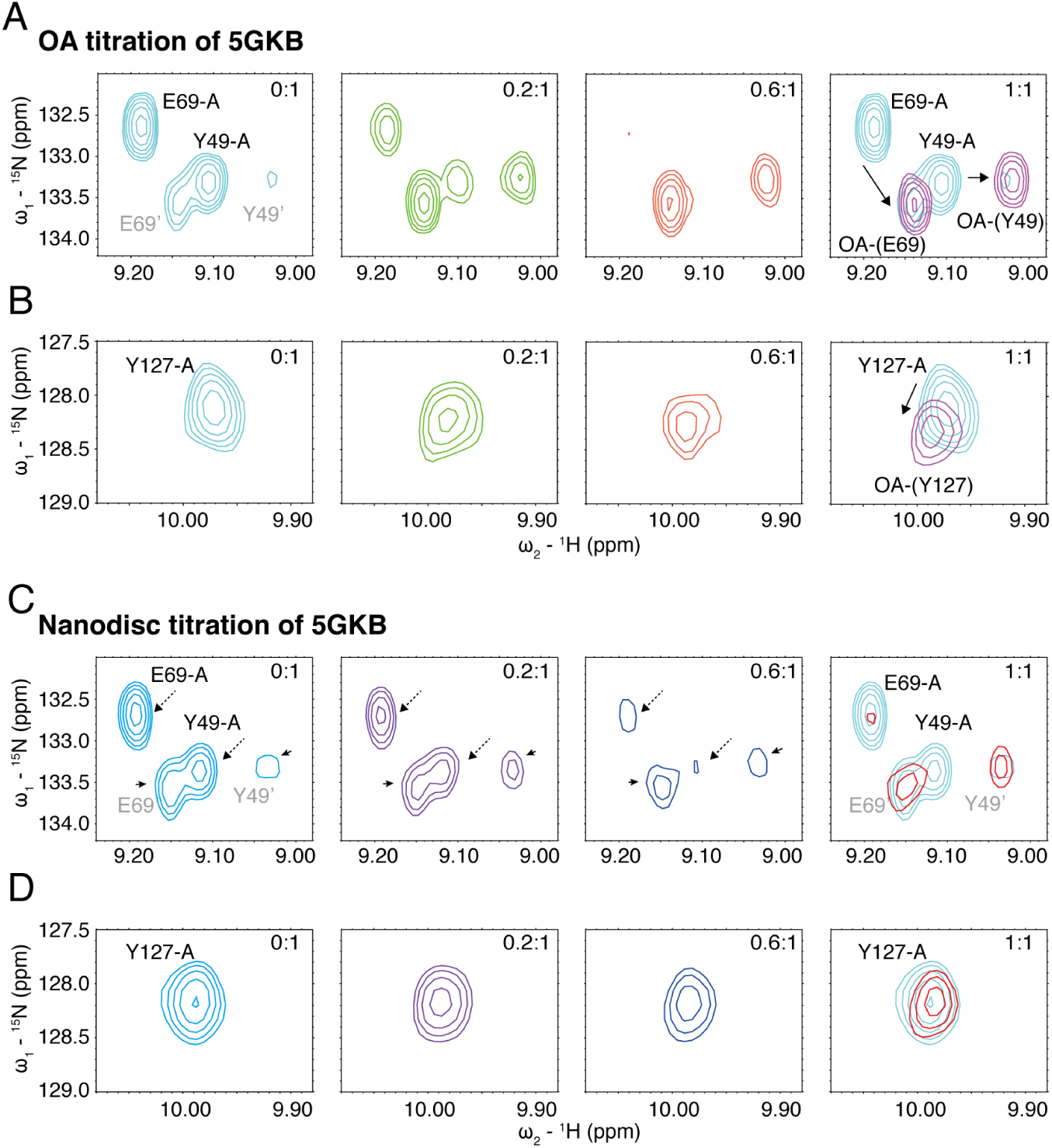
Comparison of residues in different regions of dFABP in terms of their effects on ligand binding and membrane interaction. **(A, B)** Four representative ratios used in the ligand titration of GuHCl-purified apo-dFABP are presented. The ligand OA titration for E69 and Y49, which are located near the membrane attachment (*β*E) and ligand entry (*β*B) sites, respectively, shows that conformer-A disappeared sooner during ligand binding. The ligand titration of Y127, which controls binding to the carboxylate heads of ligands, indicates that CSP occurred upon ligand binding (the arrow indicates CSP). **(C, D)** Four representative ratios used in the lipid ND titration of GuHCl-purified apo-dFABP are presented. The lipid ND titrations for E69 and Y49 show that conformer-A disappeared more slowly during lipid ND titration (dashed arrows) than during ligand titration. In contrast, the conformer-B’ of both residues was nearly unchanged (arrows) during membrane attachment. The lipid ND titration of Y127 indicates few CSP changes upon lipid interaction.

**Supplementary Fig. S8.**
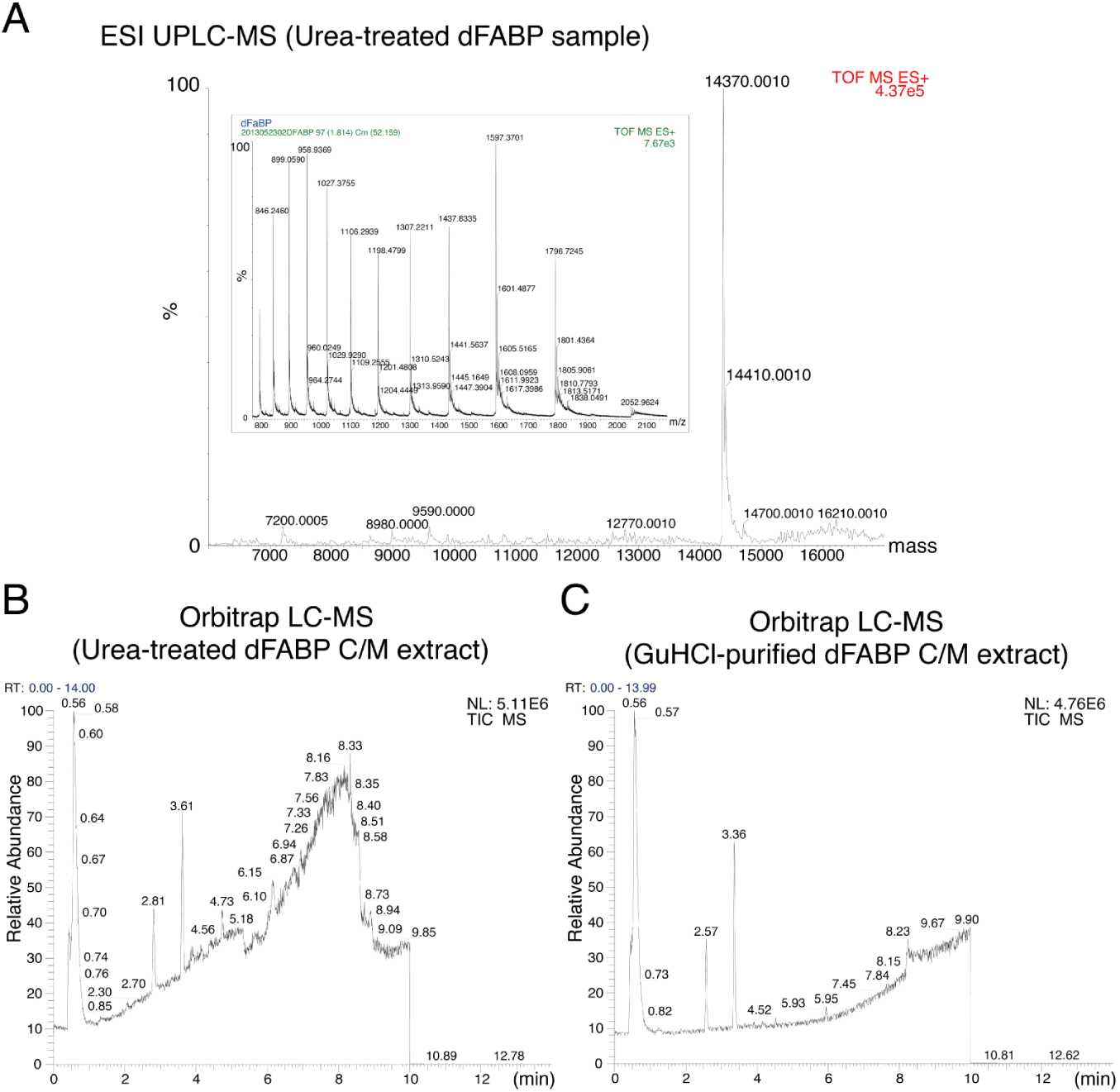
Mass spectrometry of urea and GuHCI-purified dFABP. **(A)** The theoretical molecular weight of dFABP is 14,371 Da. The urea-treated dFABP was analyzed by ESI Q-TOF UPLC-MS (Bruker, MA, USA). **(B, C)** Samples from urea and GuHCl purifications were quantified using the Bradford protein assay. Each protein solution containing 6.8 *µ*g dFABP was taken as sample for chloroform/methanol (1:1) extraction. The preceding hydrophobic compound extracts were subjected to Thermo LTQ Orbitrap liquid chromatography–mass spectrometry analysis with an ACQUITY UPLC HSS T3 column (Waters, USA)

**Supplementary Fig. S9.**
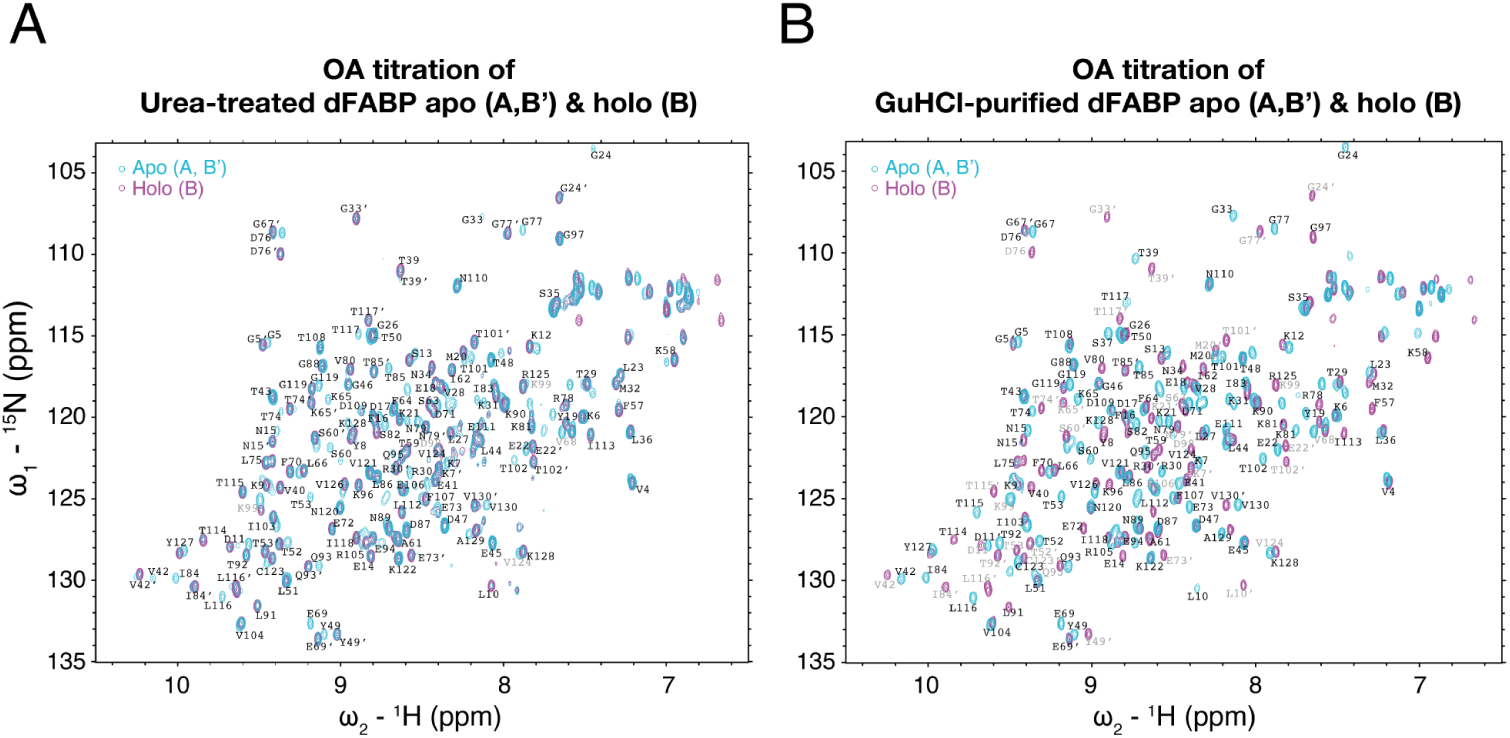
Comparison of NMR HSQC spectra in the presence and absence of ligand OA in samples with different conformational homogeneity. **(A)** The final holo form (magenta) is superimposed with the initial apo form (cyan) to compare the cross-peak dispersion of OA titration in urea-treated dFABP through HSQC. **(B)** The final holo form (magenta) is superimposed with the initial apo form (cyan) to compare the cross-peak dispersion of OA titration in GuHCl-purified dFABP through HSQC. Residues labelled with a symbol prime represent conformer-B’, and residues colored in gray exhibit a relatively low proportion compared to conformer-A.

**Supplementary Fig. S10.**
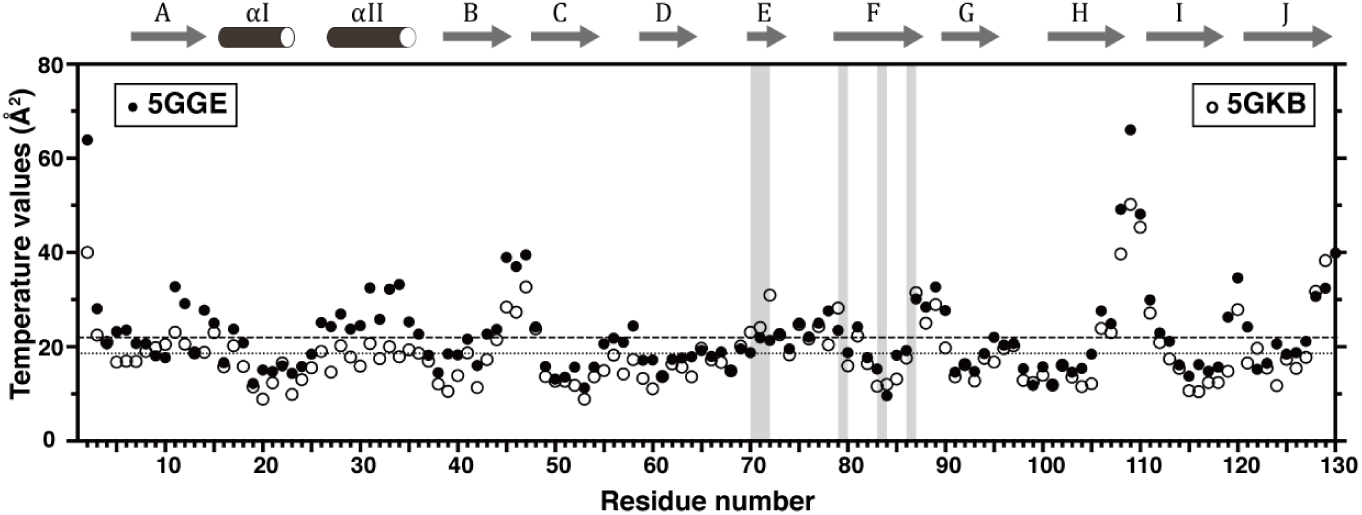
Temperature values of dFABP crystal structures. Each C*α* was plotted as a solid circle (5GGE) and an empty circle (5GKB) with the y-axis on the left and right. Higher values of residues in the gap region (*β*E and *βF*) of 5GKB are highlighted in gray stripes. Dashed and dotted lines are the average values of 5GGE and 5GKB, respectively.

**Supplementary Table S11.**
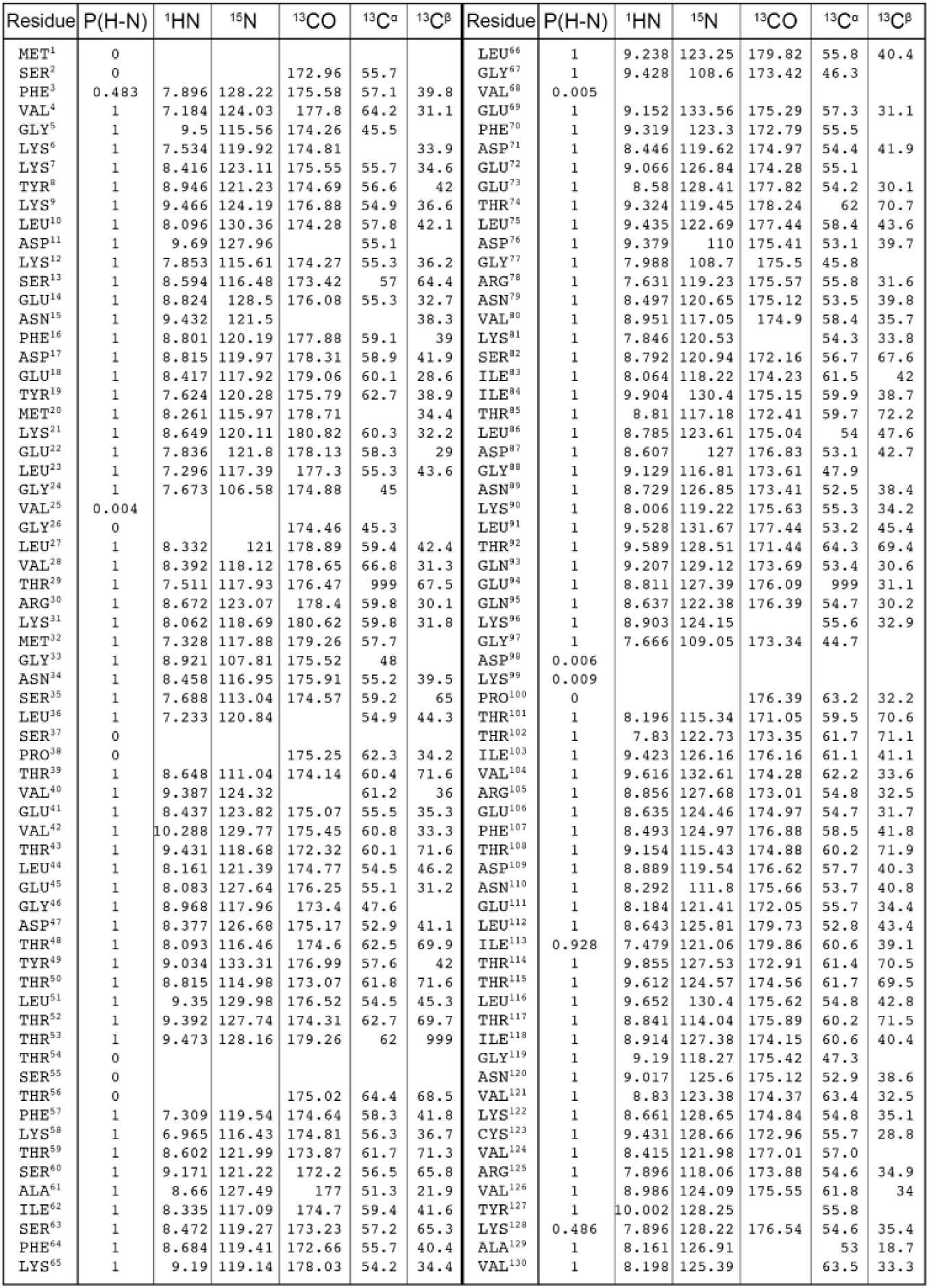
Sequential chemical shift assignments of ^15^N/^13^C OA-dFABP (BMRB ID: 27112) Autoassignment was assisted with MANI PINE server v.2.0. The MANI PINE reports were manually inspected through Sparky and subsequently removed, as summarized here. The assignments of ^1^H, ^15^N, and ^13^C chemical shifts were deposited into the BioMagResBank database under the accession number 27112.

**Supplementary Table S12.** Sequential chemical shift assignments of ^15^N/^13^C apo-dFABP (BMRB ID: 27113) The ^1^H, ^15^N, and ^13^C chemical shifts of apo-dFABP were assigned through comparison to rule out the sequential assignments of liganded OA-dFABP and deposited in the BioMagResBank database under the accession number 27113.

| Residue | P(H-N) | <sup>1</sup>HN | <sup>15</sup>N | <sup>13</sup>CO | <sup>13</sup>C<sup>α</sup> | <sup>13</sup>C<sup>β</sup> | Residue | P(H-N) | <sup>1</sup>HN | <sup>15</sup>N | <sup>13</sup>CO | <sup>13</sup>C<sup>α</sup> | <sup>13</sup>C<sup>β</sup> |
| --- | --- | --- | --- | --- | --- | --- | --- | --- | --- | --- | --- | --- | --- |
| MET<sup>1</sup> | 0 |  |  |  |  |  | LEU<sup>66</sup> | 1 | 9.169 | 123.13 | 169.21 | 53.2 |  |
| SER<sup>2</sup> | 0.987 | 7.629 | 113.18 | 174.28 | 56.6 | 62.6 | GLY<sup>67</sup> | 1 | 9.289 | 108.50 | 175.53 | 43.6 |  |
| PHE<sup>3</sup> | 0.948 | 8.371 | 119.08 | 173.45 | 54.3 | 37.3 | VAL<sup>68</sup> | 1 | 7.562 | 120.74 | 172.18 | 59.1 |  |
| VAL<sup>4</sup> | 1 | 7.109 | 123.76 | 171.13 | 61.4 | 28.6 | GLU<sup>69</sup> | 0.998 | 9.102 | 132.58 | 174.04 |  |  |
| GLY<sup>5</sup> | 1 | 9.396 | 115.33 | 174.71 | 42.9 |  | PHE<sup>70</sup> | 0.716 | 8.974 | 126.71 | 172.00 | 60.2 | 39.3 |
| LYS<sup>6</sup> | 1 | 7.437 | 119.74 | 174.11 | 52.7 | 31.4 | ASP<sup>71</sup> | 0.998 | 8.728 | 114.74 | 174.54 |  |  |
| LYS<sup>7</sup> | 1 | 8.275 | 122.65 | 171.09 | 56.4 | 36.4 | GLU<sup>72</sup> | 0.997 | 8.246 | 120.76 | 174.05 | 56.7 |  |
| TYR<sup>8</sup> | 0.525 | 8.730 | 119.79 |  |  |  | GLU<sup>73</sup> | 0.697 | 7.386 | 120.78 |  | 60.2 | 29.4 |
| LYS<sup>9</sup> | 0 |  |  |  |  |  | THR<sup>74</sup> | 0.694 | 8.119 | 115.28 |  |  |  |
| LEU<sup>10</sup> | 0.534 | 8.552 | 125.62 | 174.68 | 55.2 | 39.3 | LEU<sup>75</sup> | 0.377 | 9.000 | 118.75 | 171.58 | 55.7 |  |
| ASP<sup>11</sup> | 1 | 9.578 | 127.66 | 173.97 | 52.5 | 43.5 | ASP<sup>76</sup> | 1 | 9.333 | 109.25 | 173.62 | 50.5 |  |
| LYS<sup>12</sup> | 1 | 7.742 | 115.53 | 174.64 | 52.6 | 33.5 | GLY<sup>77</sup> | 1 | 7.812 | 108.34 | 173.41 | 43.3 |  |
| SER<sup>13</sup> | 1 | 8.476 | 116.08 | 175.53 | 54.4 | 61.7 | ARG<sup>78</sup> | 1 | 7.537 | 119.08 | 173.39 | 55.2 | 29.0 |
| GLU<sup>14</sup> | 1 | 8.754 | 128.24 | 172.87 | 52.6 | 30.1 | ASN<sup>79</sup> | 1 | 8.398 | 120.37 | 173.82 | 50.8 | 37.2 |
| ASN<sup>15</sup> | 1 | 9.357 | 121.37 | 172.89 | 51.6 |  | VAL<sup>80</sup> | 1 | 8.885 | 117.01 |  |  |  |
| PHE<sup>16</sup> | 0.995 | 8.652 | 120.31 | 171.20 | 57.0 |  | LYS<sup>81</sup> | 0.337 | 8.264 | 115.87 | 173.67 | 59.6 | 31.5 |
| ASP<sup>17</sup> | 0.917 | 8.682 | 119.82 |  | 56.1 |  | SER<sup>82</sup> | 1 | 8.560 | 111.31 | 172.02 | 57.8 |  |
| GLU<sup>18</sup> | 0 |  |  |  |  |  | ILE<sup>83</sup> | 1 | 9.766 | 127.08 | 174.98 | 58.5 |  |
| TYR<sup>19</sup> | 1 | 7.534 | 120.13 |  | 60.0 | 36.3 | ILE<sup>84</sup> | 1 | 9.928 | 129.73 | 173.81 | 57.3 | 36.0 |
| MET<sup>20</sup> | 1 | 8.163 | 115.72 | 170.32 | 57.5 | 31.8 | THR<sup>85</sup> | 1 | 8.729 | 117.00 | 176.57 | 57.0 | 69.5 |
| LYS<sup>21</sup> | 1 | 8.556 | 119.93 | 168.5 | 57.5 | 29.6 | LEU<sup>86</sup> | 1 | 8.697 | 123.47 | 173.88 | 51.3 | 44.9 |
| GLY<sup>22</sup> | 1 | 7.786 | 121.75 | 170.82 | 55.6 | 26.4 | ASP<sup>87</sup> | 1 | 8.523 | 126.75 | 172.16 | 50.5 | 40.0 |
| LEU<sup>23</sup> | 1 | 7.222 | 117.23 | 171.70 | 52.7 | 41.2 | GLY<sup>88</sup> | 1 | 9.037 | 116.64 | 175.30 | 45.3 |  |
| GLY<sup>24</sup> | 1 | 7.568 | 106.16 |  | 42.2 |  | ASN<sup>89</sup> | 1 | 8.637 | 126.62 | 175.54 | 49.8 | 35.8 |
| VAL<sup>25</sup> | 0.243 | 8.040 | 107.64 |  |  |  | LYS<sup>90</sup> | 1 | 7.922 | 118.99 | 173.32 | 52.5 | 31.5 |
| GLY<sup>26</sup> | 0.367 | 8.044 | 107.60 | 171.99 | 48.5 |  | LEU<sup>91</sup> | 1 | 9.438 | 131.36 | 172.51 | 51.9 | 42.8 |
| LEU<sup>27</sup> | 0.996 | 8.252 | 116.89 | 170.03 | 56.7 | 39.7 | THR<sup>92</sup> | 1 | 9.252 | 127.43 | 174.65 | 59.9 | 67.0 |
| VAL<sup>28</sup> | 1 | 8.300 | 117.98 | 170.27 | 64.0 | 28.7 | GLN<sup>93</sup> | 1 | 9.387 | 127.89 |  | 57.0 | 27.5 |
| THR<sup>29</sup> | 1 | 7.435 | 117.82 |  | 63.8 |  | GLU<sup>94</sup> | 0.998 | 7.959 | 118.67 |  | 58.0 |  |
| ARG<sup>30</sup> | 0.111 | 7.752 | 122.68 | 173.88 | 59.0 |  | GLN<sup>95</sup> | 1 | 9.365 | 124.19 | 172.58 | 52.1 | 27.8 |
| LYS<sup>31</sup> | 0.999 | 8.639 | 116.79 | 168.38 | 57.0 |  | LYS<sup>96</sup> | 1 | 8.832 | 124.04 | 173.10 | 52.8 | 30.3 |
| MET<sup>32</sup> | 1 | 7.256 | 117.67 | 169.59 | 55.0 |  | GLY<sup>97</sup> | 1 | 7.569 | 109.98 | 175.60 | 40.0 |  |
| GLY<sup>33</sup> | 1 | 8.942 | 107.84 | 173.32 | 45.2 |  | ASP<sup>98</sup> | 1 | 8.339 | 121.93 | 171.93 | 54.3 | 38.0 |
| ASN<sup>34</sup> | 1 | 8.371 | 117.23 | 173.10 | 52.6 | 36.8 | LYS<sup>99</sup> | 1 | 7.786 | 117.76 |  | 50.0 | 30.6 |
| SER<sup>35</sup> | 1 | 7.590 | 112.91 | 174.44 | 56.4 | 62.3 | PRO<sup>100</sup> | 0 |  |  | 172.59 | 60.5 |  |
| LEU<sup>36</sup> | 1 | 7.153 | 120.80 | 172.08 | 52.3 | 41.6 | THR<sup>101</sup> | 0.296 | 8.111 | 115.25 | 177.92 |  |  |
| SER<sup>37</sup> | 0 |  |  |  |  |  | THR<sup>102</sup> | 0.613 | 7.83 | 122.73 | 173.35 | 61.7 | 71.1 |
| PRO<sup>38</sup> | 0 |  |  | 177.48 | 61.5 |  | ILE<sup>103</sup> | 1 | 9.338 | 126.08 | 172.77 | 58.3 | 38.5 |
| THR<sup>39</sup> | 1 | 7.513 | 117.86 | 172.23 | 63.5 |  | VAL<sup>104</sup> | 1 | 9.537 | 132.46 | 173.13 | 59.6 | 31.0 |
| VAL<sup>40</sup> | 1 | 8.498 | 123.13 |  | 58.2 |  | ARG<sup>105</sup> | 0.208 | 8.540 | 124.29 | 174.00 | 52.0 | 29.1 |
| GLU<sup>41</sup> | 0.987 | 8.395 | 123.95 | 173.80 | 52.8 |  | GLU<sup>106</sup> | 1 | 8.635 | 124.46 | 174.97 | 54.7 | 31.7 |
| VAL<sup>42</sup> | 1 | 10.093 | 129.76 | 173.51 | 58.0 | 30.7 | PHE<sup>107</sup> | 1 | 8.409 | 124.77 | 172.08 | 55.7 | 39.2 |
| THR<sup>43</sup> | 1 | 9.343 | 118.57 | 176.62 | 57.3 | 68.9 | THR<sup>108</sup> | 1 | 9.059 | 115.38 | 174.03 | 57.5 | 69.1 |
| LEU<sup>44</sup> | 1 | 8.063 | 121.27 | 174.19 |  |  | ASP<sup>109</sup> | 1 | 8.805 | 119.39 | 172.34 | 55.0 | 37.6 |
| GLU<sup>45</sup> | 1 | 7.993 | 127.54 | 172.69 | 52.3 | 28.6 | ASN<sup>110</sup> | 1 | 8.206 | 111.67 | 173.72 | 50.8 | 38.2 |
| GLY<sup>46</sup> | 1 | 8.878 | 117.79 | 175.55 | 44.9 |  | GLU<sup>111</sup> | 0.529 | 9.003 | 118.75 | 173.68 | 54.5 | 28.4 |
| ASP<sup>47</sup> | 1 | 8.377 | 126.68 | 175.17 | 52.9 | 41.1 | LEU<sup>112</sup> | 0.996 | 9.225 | 123.22 | 176.20 |  |  |
| THR<sup>48</sup> | 1 | 7.974 | 116.11 | 174.33 | 59.8 | 67.2 | ILE<sup>113</sup> | 0.682 | 8.338 | 119.23 | 177.28 | 56.8 |  |
| TYR<sup>49</sup> | 1 | 9.019 | 133.10 |  | 54.9 |  | THR<sup>114</sup> | 0.623 | 7.874 | 122.38 | 176.09 | 58.6 | 67.8 |
| THR<sup>50</sup> | 0 |  |  |  |  |  | THR<sup>115</sup> | 0.656 | 9.525 | 124.40 | 174.42 | 59.0 | 66.8 |
| LEU<sup>51</sup> | 0.627 | 8.887 | 120.25 | 176.17 |  |  | LEU<sup>116</sup> | 0.971 | 9.595 | 130.19 | 173.39 | 52.0 | 40.1 |
| THR<sup>52</sup> | 0.743 | 9.283 | 119.43 | 171.07 |  |  | THR<sup>117</sup> | 1 | 8.778 | 113.83 | 173.48 | 57.4 | 68.8 |
| THR<sup>53</sup> | 0.792 | 9.219 | 119.35 |  |  | 68.0 | ILE<sup>118</sup> | 1 | 8.797 | 127.07 | 174.80 | 57.6 |  |
| THR<sup>54</sup> | 0.273 | 9.524 | 124.39 | 175.01 | 59.3 | 67.1 | GLY<sup>119</sup> | 1 | 9.060 | 117.79 | 173.93 | 43.4 |  |
| SER<sup>55</sup> | 0.416 | 9.385 | 122.63 |  | 53.8 | 63.1 | ASN<sup>120</sup> | 0.911 | 9.615 | 119.63 | 173.82 | 50.2 | 36.0 |
| THR<sup>56</sup> | 0.658 | 8.576 | 127.40 | 173.88 | 61.6 | 65.7 | VAL<sup>121</sup> | 1 | 8.748 | 123.24 | 174.58 | 60.7 | 29.9 |
| PHE<sup>57</sup> | 1 | 7.220 | 119.06 | 174.21 | 55.5 | 39.0 | LYS<sup>122</sup> | 1 | 8.570 | 128.45 |  | 52.2 | 32.4 |
| LYS<sup>58</sup> | 1 | 6.924 | 116.38 | 174.16 | 53.6 | 33.8 | CYS<sup>123</sup> | 1 | 9.367 | 128.63 | 176.00 | 53.0 | 26.1 |
| THR<sup>59</sup> | 1 | 8.532 | 122.04 |  | 59.0 | 68.5 | VAL<sup>124</sup> | 1 | 7.816 | 128.11 | 177.40 | 58.7 |  |
| SER<sup>60</sup> | 0.182 | 8.127 | 116.17 | 177.50 | 56.3 | 64.3 | ARG<sup>125</sup> | 1 | 9.531 | 130.47 |  |  |  |
| ALA<sup>61</sup> | 1 | 9.077 | 128.98 | 175.31 | 50.7 |  | VAL<sup>126</sup> | 1 | 8.908 | 124.26 | 174.17 | 59.0 |  |
| ILE<sup>62</sup> | 1 | 8.736 | 127.27 | 175.12 | 59.3 |  | TYR<sup>127</sup> | 0.996 | 9.890 | 128.01 | 173.60 |  |  |
| SER<sup>63</sup> | 0.999 | 8.999 | 121.71 | 176.83 | 54.0 | 65.1 | LYS<sup>128</sup> | 0.996 | 8.857 | 120.66 | 172.46 | 52.1 |  |
| PHE<sup>64</sup> | 0.998 | 7.977 | 118.04 | 174.68 | 58.7 | 39.3 | ALA<sup>129</sup> | 1 | 8.126 | 126.97 | 171.52 | 50.4 | 16.1 |
| LYS<sup>65</sup> | 0.900 | 9.833 | 130.26 |  |  | 31.8 | VAL<sup>130</sup> | 1 | 8.038 | 125.17 |  |  |  |

**Supplementary Table S13.** Structure statistics of crystallographic data.

| Dataset | Monomer | Dataset | Monomer |
| --- | --- | --- | --- |
| Protein sample | dFABP<sub>2-130</sub> (5GGE) | Protein sample | dFABP<sub>2-130</sub> (5GKB) |
| Beamline | NSRRC_BL13B1 | Beamline | NSRRC_BL13C1 |
| Wavelength (Å) | 0.97622 | Wavelength (Å) | 0.97622 |
| Resolution (Å) | 27.05-1.86 (1.93-1.86) | Resolution (Å) | 30.00-2.04 (2.11-2.04) |
| Space group | I422 | Space group | I422 |
| *Unit cell dimensions* |  | *Unit cell dimensions* |  |
| a, b, c (Å) | 61.70, 61.70, 137.32 | a, b, c (Å) | 61.60, 61.60, 137.06 |
| a, b, g (°) | 90 | a, b, g (°) | 90 |
| Unique reflections | 11518 | Unique reflections | 8811 |
| Total observations | 79527 | Total observations | 81557 |
| Redundancy | 6.9 (6.2) | Redundancy | 9.3 (9.1) |
| Completeness (%) | 99.5 | Completeness (%) | 99.9 |
| Mean I/S (I) | 27.277 (3.2) | Mean I/S (I) | 26.900 (6.1) |
| Solvent content (%) | 41.6 | Solvent content (%) | 45.1 |
| R merge | 0.068 | R merge | 0.100 |
| Refinement |  | Refinement |  |
| Resolution range (Å) | 27.052-1.861 | Resolution range (Å) | 28.091-2.039 |
| No. of reflections | 26.98-1.86 (1.93-1.86) | No. of reflections | 28.091-2.04 (2.11-2.04) |
| No. of protein atoms | 2229 | No. of protein atoms | 2146 |
| No. of water molecules | 139 | No. of water molecules | 128 |
| Average B-factors (Å<sup>2</sup>) | 30 | Average B-factors (Å<sup>2</sup>) | 24 |
| R work/R free | 0.179/0.246 | R work/R free | 0.186/0.241 |
| *Ramachandran plot statistics (%)* |  | *Ramachandran plot statistics (%)* |  |
| Most favored regions | 99.21 | Most favored regions | 98.41 |
| Additional allowed regions | 0.85 | Additional allowed regions | 2.59 |
| Generously allowed regions | 0.79 | Generously allowed regions | 1.59 |
| Disallowed regions | 0.00 | Disallowed regions | 0.00 |
| Rmsd bond length (Å) | 0.56 | Rmsd bond length (Å) | 0.77 |
| Rmsd bond angle (°) | 0.69 | Rmsd bond angle (°) | 0.83 |

**Supplementary Table S14.**
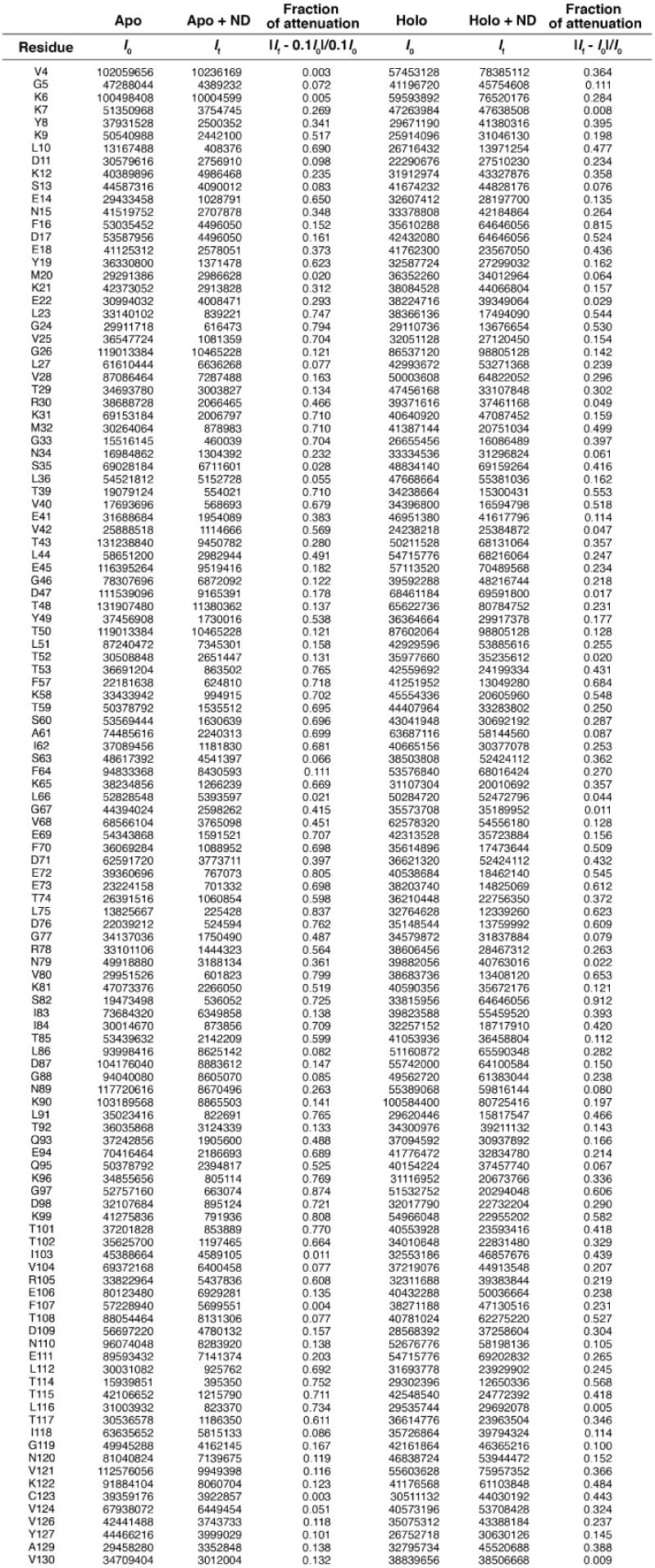
Intensity changes during lipid ND titration.

